# Mechanical phenotyping of acute promyelocytic leukemia reveals unique biomechanical responses in retinoic acid-resistant populations

**DOI:** 10.1101/2021.04.25.441378

**Authors:** Brian Li, Annie Maslan, Aaron M. Streets, Lydia L. Sohn

## Abstract

While all-trans retinoic acid (ATRA) is an essential therapy in the treatment of acute promyelocytic leukemia (APL), an aggressive subtype of acute myeloid leukemia, nearly 20% of APL patients are resistant to ATRA. As no biomarkers for ATRA resistance yet exist, we investigated whether cell mechanics could be associated with this pathological phenotype. Using mechano-node-pore sensing, a single-cell mechanical phenotyping platform, and patient-derived APL cell lines, NB4 (ATRA-sensitive) and AP-1060 (ATRA-resistant), we discovered that ATRA-resistant APL cells are less mechanically pliable. By investigating how different subcellular components of APL cells contribute to whole-cell mechanical phenotype, we determined that nuclear mechanics strongly influence an APL cell’s mechanical response. By arresting APL cells in S-phase or M-phase in the cell cycle, we found cell pliability to be inversely related to DNA content. In addition to DNA content affecting cell pliability, we observed that chromatin condensation also affects nuclear mechanics: decondensing chromatin with trichostatin A is especially effective in softening ATRA-resistant APL cells. RNA-Seq allowed us to compare the transcriptomic differences between ATRA-resistant and ATRA-responsive APL cells and highlighted gene expression changes that could be associated with mechanical changes. Overall, we demonstrate the potential of “physical” biomarkers in identifying APL resistance.

## Introduction

Increasingly, physical and mechanical properties such as mass growth rate and deformability, respectively, are being explored as possible biomarkers of cancer cells (1–6). While cells from solid tumors have been the primary focus of these studies, cells from liquid tumors have also been investigated, albeit to a lesser extent. Work using atomic-force microscopy and optical stretchers have shown that deformability and mechanical compliance are important in the pathological dissemination of leukemic cells throughout the circulatory system (7–11). More recently, suspended microchannel resonators studies have correlated physical properties (*e*.*g*., cell mass and growth rate, of leukemias and lymphomas to important pathological processes such as drug sensitivity or proliferation (12, 13), and investigated some of the biophysical factors that influence cell mechanical properties, such actin remodeling and cell cycle phase (14). Despite these investigations, the relevance of the physical and mechanical attributes of liquid tumor cells to sensitivity to therapy has yet to be examined (7, 15).

Here, we assess the cell mechanics of Acute Promyelocytic Leukemia (APL), an acute myeloid leukemia subtype characterized by a fusion gene between PML (promyelocytic leukemia protein) and RARA (retinoic acid receptor alpha) (16). In APL, the PML-RARA fusion gene occurs in granulocytic-precursor cells known as promyelocytes, halting their differentiation and causing rapid proliferation of the immature blasts. Because of the PML-RARA fusion gene, the standard of care for APL patients is to administer all-trans retinoic acid (ATRA), which induces differentiation of the promyelocytes and, in turn, resolves the acute phase of the disease (17–20). This acute phase can rapidly progress to mortality within a week if left untreated. Early deaths due to disseminated intravascular coagulopathy are of especially great risk in *ATRA-resistant* cases, which comprise nearly 20% of all APL cases (21–24). Although APL has been well-characterized biochemically using cDNA microarrays and RT-PCR (25–28), there has only been limited characterization of the cell mechanics in APL. Specifically, Lautenschläger *et al*. showed that the APL cell line, NB4, becomes more deformable with ATRA treatment as it differentiates (9). However, beyond this particular study, there have been no investigations of the mechanical responses of ATRA-resistant APL to our knowledge.

We used mechano-node-pore sensing (mechano-NPS) (5), a microfluidic platform that utilizes a narrow channel to deform cells and measure their mechanical properties, to investigate ATRA-sensitive and ATRA-resistant APL cells. Through our mechano-NPS measurements, we determined that ATRA-resistant APL cells have a unique mechanical phenotype that is influenced by the organization of various structural subcellular components. Furthermore, using chemical cell-cycle arrest and perturbation of chromatin accessibility, we evaluated how both major and minor changes in nuclear structure, respectively, affect APL mechanical phenotypes. Through these studies, we found that ATRA-resistant APL cells are uniquely affected by pharmacological chromatin decondensation. By performing RNA-Seq analysis of ATRA-resistant and ATRA-sensitive cells, we have obtained a whole-transcriptome perspective of how gene expression changes in response to ATRA. This work establishes a resource for further studies of ATRA resistance in APL. Overall, our studies show that ATRA-resistance in APL corresponds to unique biomechanical responses.

## Results

### Single-cell mechanical phenotyping of acute promyelocytic leukemia cells

Mechano-NPS (5) is an electronic method for mechanophenotyping cells that involves measuring the current across a microfluidic channel segmented by widened “nodes” (Fig. 1A, B; see Materials and Methods for fabrication and platform operation details). The width of one segment is narrower than a cell diameter (hereafter referred to as the “contraction” segment), and cells must squeeze through this segment in order to traverse the entire channel (Fig. 1C). Wider “recovery” segments immediately following the contraction segment allow the cell to recover from deformation. When a cell transits the channel, a unique current pulse is measured. This current pulse is comprised of subpulses that correspond to the cell traversing specific mechano-NPS segments (Fig. 1D) (see Materials and Methods and Supplementary Note 1). It is these subpulses from which physical and mechanical properties of the cell can be extracted. Specifically, the magnitude and duration of the subpulse prior to the cell entering the contraction segment reflect the cell size and velocity, respectively (Supplementary Movie 1). The dramatically large subpulse corresponds to the cell entering the contraction segment (Supplementary Movie 2), and its duration provides information on the cell’s deformability (*i*.*e*., a more deformable cell will transit the contraction segment faster than a stiffer cell). Finally, the series of subpulses produced by the cell transiting the series of node-pores following the contraction segment tracks the cell’s recovery from deformation. As the cell relaxes back to its original size and shape, the magnitude of the subpulses return to that of the initial subpulse, i.e., the one caused by the cell initially entering the first initial node-pore segment (Supplementary Movie 3).

**Figure 1:**
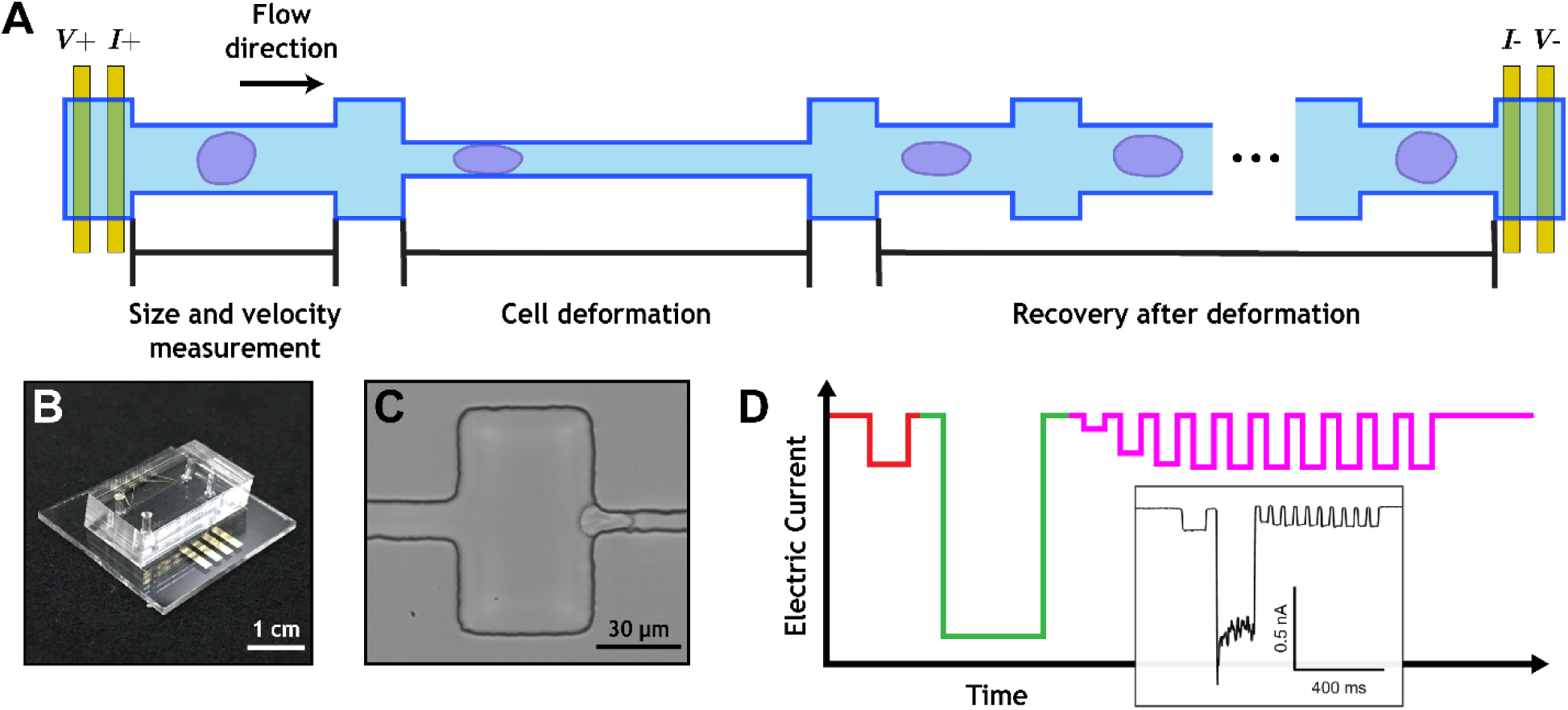
Overview of mechano-node-pore sensing (mechano-NPS) platform. **A**. Top view schematic of mechano-NPS microchannel. A four-terminal measurement is used to measure the current across the microchannel. Cell size (*d*_0_) and velocity (*v*_0_) are measured prior to cell deformation. During deformation, the velocity of a cell is dependent on the cell’s elastic modulus. After it leaves the contraction channel, the cell recovers from an ellipsoid shape to a sphere. **B**. Photograph of a mechano-NPS platform, with the PDMS mold embedded with the mechano-NPS channel and bonded to a glass substrate with prefabricated metal electrodes. **C**. High-speed image of an NB4 cell being deformed in transit through the contraction channel. **D**. Expected current pulse caused by a cell transiting the entire mechano-NPS channel. The current pulse consists of subpulses that reflect the three main regions of the microchannel. (1) In the initial cell sizing segment (red), the free diameter and velocity of the cell are measured by the subpulse amplitude and duration, respectively. (2) In the contraction segment (green), the constricted velocity of the cell is measured by the subpulse duration, where more deformable cells transit this segment faster. (3) In the 10 cell recovery segments (magenta), cells recover from a deformed ellipsoid shape to a sphere, which is reflected in the gradual increase in subpulse amplitude over time. The inset shows an actual current pulse produced by an NB4 cell transiting a 6 mm long mechano-NPS channel at an inlet pressure of 80 mbar.

A cell’s mechanical response is dependent on the degree of applied strain, ε:

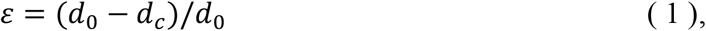

where *d*_0_ is the cell’s initial diameter and *d*_c_ is the cell’s deformed diameter. For our studies, ε ≈ 0.35 in the contraction segment. To describe a cell’s deformability, we use the whole-cell deformability index (*wCDI*), a dimensionless number that relates a cell’s size and velocity through the contraction segment (5):

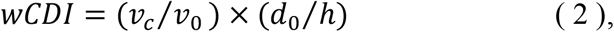

where *v*_c_ is the cell velocity through the contraction segment, *v*_0_ is the cell’s velocity as it enters the contraction segment, *d*_0_ is the cell’s initial diameter, and *h* is the microfluidic channel height. As was previously established (5), *wCDI* is inversely proportional to the elastic modulus: stiffer cells have a smaller *wCDI* than softer cells.

As cells are intrinsically viscoelastic materials, the time-dependent response to changes in applied stress is a critical aspect of cell mechanical phenotype. To measure cell viscoelastic behavior, we use the recovery segments in a mechano-NPS channel to quantify how cells recover from deformation. For the experiments we report here, we designed a device with 10 recovery channel segments, which provided sufficient information to calculate the time constant of a cell’s recovery using linear least squares regression for a Kelvin-Voigt model (Supplementary Note 1). This recovery time ranges from 20 ms to 800 ms, where longer recovery times are indicative of more viscous behavior (Supplementary Fig. S1).

### ATRA-resistant APL cells are phenotypically and mechanically distinct

We investigated two human APL cell lines: NB4, derived from an APL patient who retained the characteristic sensitivity to ATRA differentiation; and AP-1060, derived from an APL patient who displayed ATRA resistance (29–31). We cultured NB4 and AP-1060 cells in media supplemented with ATRA in concentrations ranging from 10 nM to 10 µM and subsequently evaluated their propensity to differentiate (see Materials and Methods). Using flow cytometry to measure expression of CD11b and CD18, two integrins expressed by mature innate immune cells, we confirmed that while NB4 cells respond to ATRA, AP-1060 cells were largely resistant (Supplementary Fig. S2, S3) (32).

The differences between AP-1060 and NB4 cells extend well beyond their biochemical response to ATRA. Upon mechanophenotyping these cells, we found that untreated AP-1060 cells have a significantly lower *wCDI* (*p* < 0.0001) than that of untreated NB4 cells, indicating that they are stiffer (Fig. 2A). After treating both cell lines with two doses of 1 µM ATRA over the course of 4 days (see Materials and Methods), we observed that only NB4 cells showed a significant change in their *wCDI* (*p* < 0.0001) and became more deformable (Fig. 2A). Although the ATRA dose we chose did lead to a limited degree of maturation in AP-1060 cells (Supplementary Fig. S3), there was still no significant change in their overall deformability.

**Figure 2:**
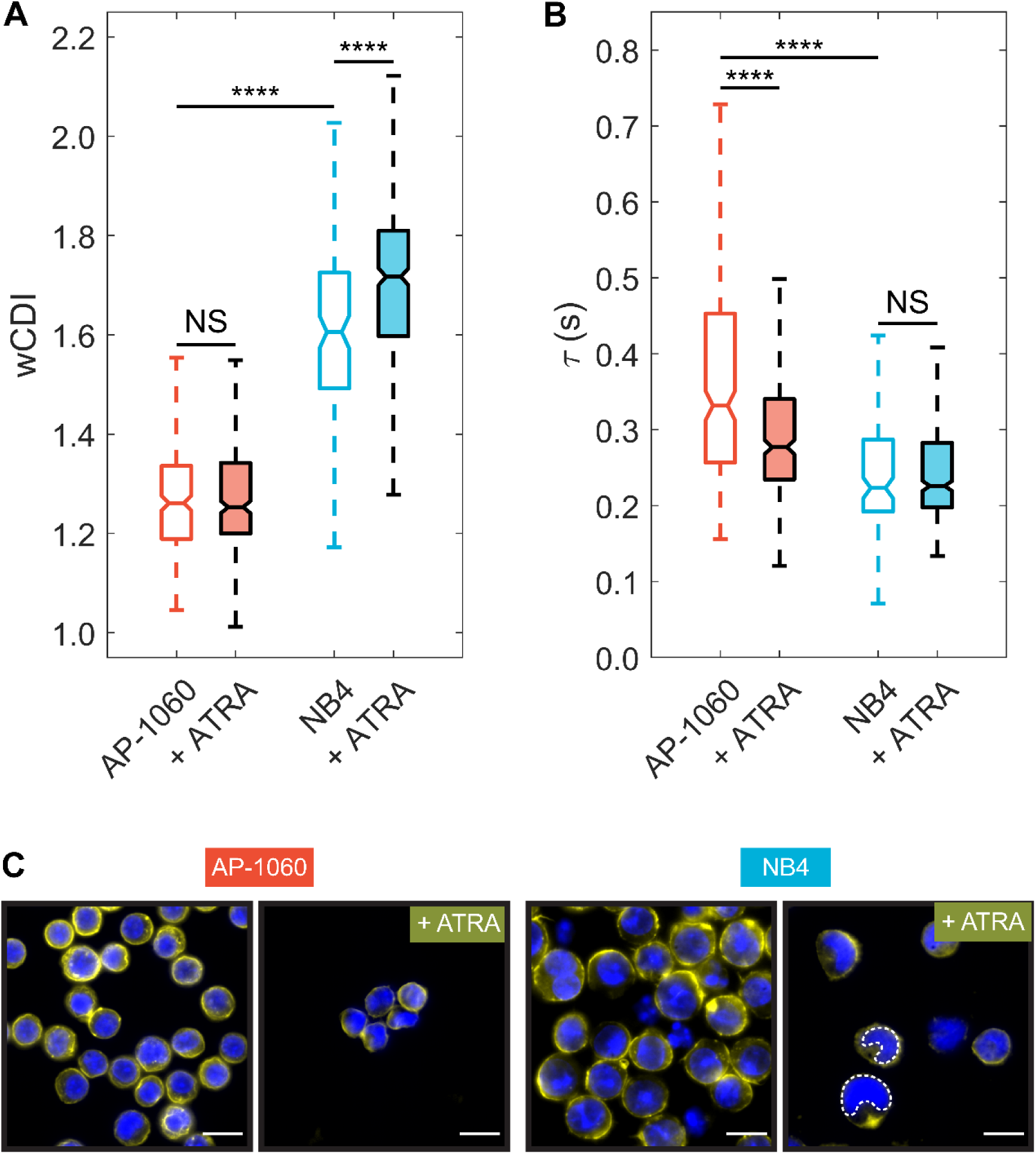
APL cell responses to all-trans retinoic acid (ATRA). **A, B**. Box plots of whole cell deformability index (*wCDI*) (**A**) and recovery times (**B**) for AP-1060 and NB4 cells before and after treatment with ATRA. Notches represent 95% confidence intervals for the true median of each distribution. *n* ≥ 124 for each distribution across 3 different devices. \*\*\*\**p* < 0.0001, NS corresponds to no significance; statistical significance was determined by a Tukey test for multiple comparisons. **C**. Fluorescence images of AP-1060 and NB4 cells before and after treatment with ATRA. Cell nuclei stained with Hoescht 33342 (blue); F-actin stained with rhodamine phalloidin (yellow). Lobulated (U-shaped) nuclei in ATRA-treated NB4 cells are highlighted with a white dashed outline. Scale bar = 15 µm.

As with their *wCDI*s, we observed differences between the apparent viscosities of AP-1060 and NB4 cells via their recovery times. AP-1060 cells took far longer to recover from deformation than NB4 cells (*p* < 0.0001) and were therefore more viscous (Fig. 2B). Unexpectedly, while ATRA-treated AP-1060 recovered significantly faster from deformation than untreated AP-1060 cells (*p* < 0.0001), we observe no significant difference in the recovery times of ATRA-treated and untreated NB4 cells (*p* = 0.99). Although AP1060 cells are resistant to ATRA, our analysis of CD11b and CD18 expression showed that they are not completely unresponsive to ATRA when introduced to a high concentration (Supplementary Fig. S3). Interestingly, we found that this partial response is also associated with a partial response in mechanical phenotype where changes in recovery time, but not *wCDI*, occur. Ultimately, however, our results show the differential response to ATRA between AP-1060 and NB4 cells affects not only protein expression but also mechanical phenotype, where the two cell lines had different responses in the elastic (*wCDI*) and viscoelastic (recovery time) mechanical regimes.

Because the spatial organization and reorganization of the nucleus constitute a critical part of cell mechanics (33–35), we employed fluorescence microscopy to image the stained nuclei of APL cells before and after ATRA treatment (Fig. 2C; see Materials and Methods). We observed lobulation in NB4 nuclei (recognizable by the U-shaped nuclei) after ATRA treatment, indicating that differentiation in promyelocytes was progressing (Supplementary Fig. S4) (36). In contrast, we did not observe such lobulation in ATRA-treated AP-1060 nuclei, which is consistent with AP1060 cells’ resistance to differentiate. This nuclear shape requires downregulation of lamin A, which itself increases nuclear and whole-cell deformability (37, 38). As such, the lobulation of NB4 nuclei may contribute to their softer mechanical phenotype.

In addition to the nucleus, we examined the F-actin network, which provides mechanical support for the cell and its organelles and drives cell motility (39–43). Although there were no major visual differences in the F-actin network between cell types or before and after ATRA treatment (Supplementary Fig. S5), we sought to determine how F-actin might affect cell mechanical properties. We destabilized F-actin with Latrunculin A (LatA) (Materials and Methods) and confirmed the disruption of cortex-localized F-actin in both AP-1060 and NB4 cells (Supplementary Fig. S6). Upon mechano-NPS screening of LatA-treated APL cells, we observed no significant differences in the deformability of AP-1060 (*p* = 0.61) or NB4 (*p* = 0.19) cells (Supplementary Fig. S7A). However, the recovery times for both cells with disrupted actin cortices were significantly faster than those of untreated cells (AP-1060, *p* < 0.0001; NB4, *p* = 0.0027) (Supplementary Fig. S7B), with differences more pronounced with AP-1060 cells than with NB4 cells (125 ms vs. 27 ms faster median recovery, respectively). Overall, however, since ATRA affected both the elastic and viscoelastic regimes of APL cells and LatA only affected cell viscoelasticity, we conclude that the mechanical changes in ATRA-treated APL cells are not solely due to any remodeling of the F-actin network that may arise after ATRA treatment.

### APL single-cell mechanical phenotypes vary with DNA content

Because DNA content and nuclear volume change throughout a cell’s life cycle, cell-cycle phase is also an important contributor to whole-cell mechanical properties (45–47). We demonstrated this by performing a series of experiments in which we mechanically phenotyped cells that were synchronized either in M- or S-phase (high DNA content or low DNA content, respectively) Specifically, we treated cells with colcemid (a mitotic inhibitor) to arrest them in M-phase before mechanophenotyping (Fig. 3A; Materials and Methods). Both NB4 and AP-1060 cells showed a significant increase in stiffness (*p* < 0.0001), where NB4 stiffening was more pronounced (Fig. 3B). While colcemid-treated NB4 cells also had significantly slower recovery times compared to untreated cells (*p* < 0.0001) and therefore were more viscous, colcemid-treated AP-1060 cells did not (*p* = 0.93) (Fig. 3C).

**Figure 3:**
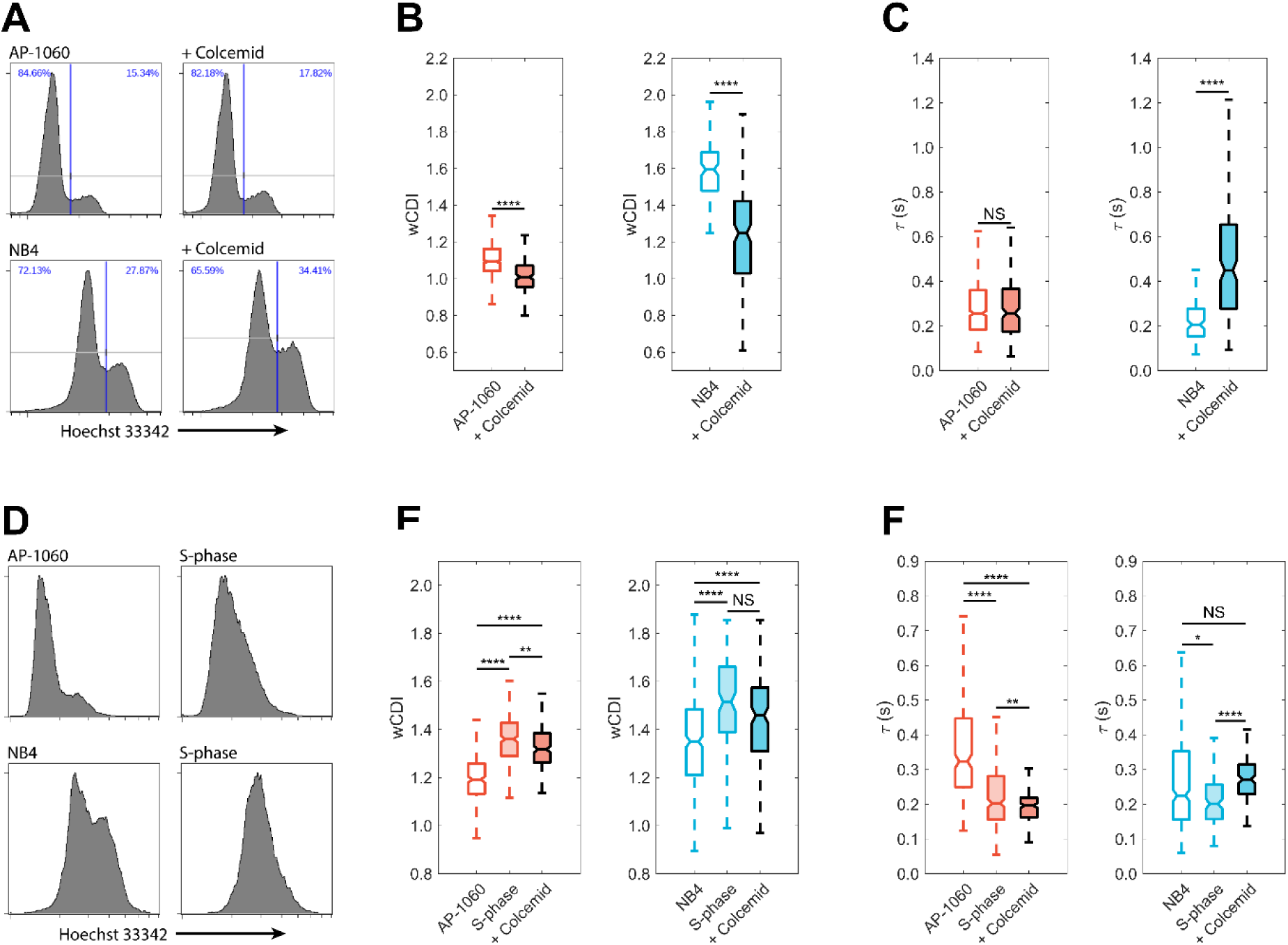
Mechanical phenotypes of APL cells are strongly influenced by cell-cycle phase. **A**. Histograms of DNA content in AP-1060 and NB4 cells before and after colcemid treatment, as measured by flow cytometry. Colcemid treatment causes small increases in cell counts staining high for DNA, indicating an increased mitotic index. **B, C**. Box plots of *wCDI* (**B**) and recovery times (**C**) for AP-1060 and NB4 cells before and after treatment with colcemid. Notches represent 95% confidence intervals for the true median of each distribution. *n* ≥ 123 for each distribution across 3 different devices. \*\*\*\**p* < 0.0001, NS no significance; statistical significance was determined by two-sample Student’s *t*-tests. **D**. Histograms of DNA content in AP-1060 and NB4 cells before and after synchronization at S-phase with a double thymidine block as measured by flow cytometry. Cell cycle synchronization reduces the bimodal distribution to a unimodal distribution, representing a decreased mitotic index. **E, F**. Box plots of *wCDI* distributions (**E**) and recovery times (**F**) for AP-1060 and NB4 cells before and after double thymidine block. Notches represent 95% confidence intervals for the true median of each distribution. *n* ≥ 122 for each distribution across 3 different devices. \**p* < 0.05, \*\*\*\**p* < 0.0001, NS no significance; statistical significance was determined by two-sample Student’s *t*-tests.

To determine whether low DNA content would produce the opposite effect, *i*.*e*. increased deformability and lower viscosity, we arrested AP-1060 and NB4 cells in S-phase using a double thymidine block (49) and subsequently performed mechanophenotyping (Fig. 3D). In contrast to when cells were in M-phase, cells in S-phase were significantly more deformable than untreated cells (*p* < 0.0001) (Fig. 3E). Moreover, compared to untreated cells, both NB4 and AP-1060 cells recovered from deformation significantly faster after the double thymidine block (NB4, *p* = 0.0105, AP-1060, *p* < 0.0001) (Fig. 3F). Thus, changes in DNA content can have opposite effects on cell mechanical properties, depending on whether DNA content increased or decreased.

As a control, we additionally assessed if chemically induced cell cycle arrest can affect mechanical phenotype by itself, without affecting DNA content. To test this, we again arrested AP-1060 and NB4 cells in S-phase via a double thymidine block and subsequently treated them with colcemid. As cells arrested in S-phase will not progress through the cell cycle, the colcemid treatment should have no effect on DNA content, and therefore, any mechanical differences due to this particular treatment would not be due to DNA content. We found that while it did not significantly affect the *wCDI* of NB4 cells (*p* = 0.39), colcemid significantly decreased the *wCDI* of AP-1060 (*p* < 0.01) (Fig. 3E), albeit to a lesser degree compared to M-phase arrested AP-1060 cells (Fig. 3B). Colcemid treatment of S-phase cells also significantly increased the recovery time of both AP-1060 (*p* < 0.01) and NB4 (*p* < 0.0001) (Fig. 3F). While we found that colcemid indeed affects the mechanical phenotype of AP-1060 and NB4 cells independently of DNA content, these effects are far less significant and dramatic than the cell stiffening we observed when cells are arrested in M-phase. Indeed, when we sorted DNA-stained NB4 cells based on DNA content (see Materials and Methods) and subsequently mechanophenotyped them, we observed that low-DNA content cells had a greater *wCDI* and faster recovery time compared to high-DNA content cells (Supplementary Fig. S8).

### Influences of histone acetylation on APL cell mechanical phenotype

We probed how cellular mechanical phenotype is affected by the mechanism by which ATRA affects chromatin accessibility to achieve differentiation of APL cells. Specifically, we evaluated how chromatin decondensation affects whole-cell mechanics. Previous studies have shown that chromatin accessibility is modulated by histone acetylation downstream of retinoid signaling pathways and that histone decondensation can soften isolated nuclei (18, 50–53). To promote chromatin decondensation within our two APL cell lines, we inhibited histone deacetylase (HDAC) activity with trichostatin A (TSA) (see Materials and Methods). To measure primarily the mechanical properties of the nucleus and the effect of TSA, we destabilized actin cortices and arrested cells in S-phase by treating AP-1060 and NB4 cells with LatA and a double thymidine block, respectively (see Materials and Methods). We then treated the cells with TSA or ATRA and subsequently mechanophenotyped them. As a control, we mechanophenotyped synchronized and LatA-treated cells without further perturbations.

TSA treatment significantly increased *wCDI* in both AP-1060 (*p* < 0.0001) and NB4 (*p* = 0.0014) cells, suggesting that chromatin decondensation indeed causes nuclear softening (Fig. 4A). Unexpectedly, ATRA treatment of actin-destabilized and S-phase arrested AP-1060 cells resulted in a dramatic decrease in *wCDI* (*p* < 0.0001), opposite to what we observed in earlier experiments when cells were simply treated with ATRA (Fig. 2A). In contrast, actin-destabilized and S-phase arrested NB4 cells were more deformable after ATRA treatment (*p* < 0.0001) (Fig. 4A, right), agreeing with the results of our earlier experiments when these cells were only treated with ATRA (Fig. 2A). Recovery time responses were similarly nuanced: TSA significantly decreased recovery times in AP-1060 (*p* = 0.0095) but not NB4 (*p* = 0.13) cells (Fig. 4B).; ATRA-treated AP-1060 cells showed no significant difference in recovery time (*p* = 0.15); but NB4 cells took significantly longer to recover (*p* = 0.0030).

**Figure 4:**
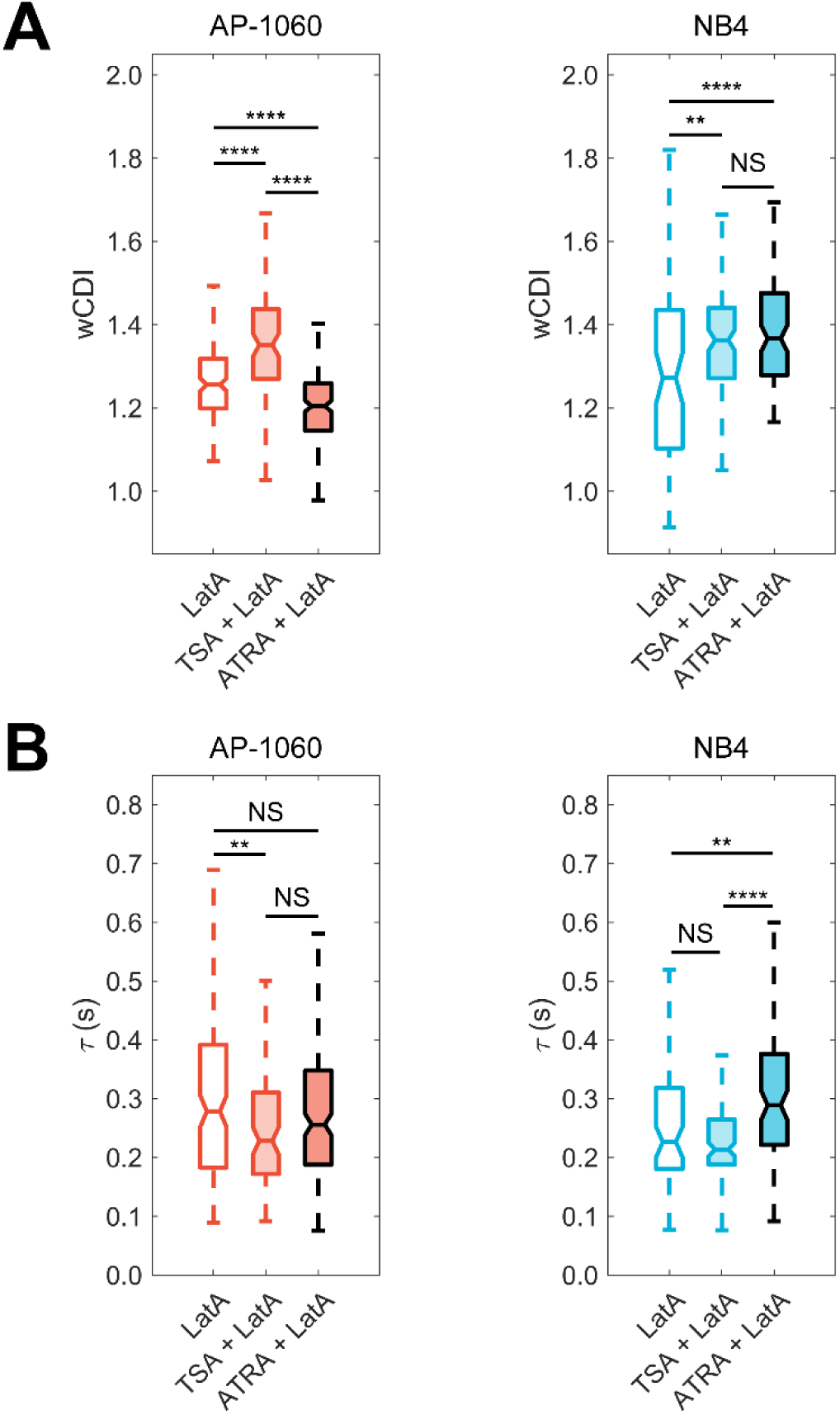
Mechanical phenotype of ATRA-resistant APL nuclei is sensitive to histone acetylation. **A**. Box plots of wCDI for S-phase synchronized AP-1060 (left) and NB4 (right) with actin cortices destabilized with Latrunculin A. Cells were treated with TSA to promote accumulation of decondensed chromatin or ATRA to induce differentiation. For AP-1060, *n* ≥ 84 for each distribution across 3 different devices. For NB4, *n* ≥ 75 for each distribution across 3 different devices. \*\**p* < 0.01, \*\*\*\**p* < 0.0001, NS no significance; statistical significance was determined by a Tukey test for multiple comparisons. **B**. Box plots of recovery times for AP-1060 (left) and NB4 (right) treated with Latrunculin A and ATRA or TSA. For AP-1060, *n* ≥ 84 for each distribution across 3 different devices. For NB4, *n* ≥ 75 for each distribution across 3 different devices. \*\**p* < 0.01, \*\*\*\**p* < 0.0001; statistical significance was determined by a Tukey test for multiple comparisons.

Overall, these measurements highlight the complexity of measuring whole-cell deformability, where the effects of certain perturbations (i.e., ATRA) on cell stiffness can change depending on the state of other subcellular components (i.e., F-actin, cell cycle). For actin-destabilized and S-phase arrested cells, TSA had a more consistent softening effect than ATRA across both cell types and mechanical parameters, most likely because TSA causes chromatin decondensation. While nuclear softening is expected, TSA treatment may also modulate the expression of genes that could affect single-cell mechanical phenotype (54, 55).

### Characterizing transcriptomic changes in APL cells with RNA-Seq

Studies involving cDNA microarrays have identified patterns of differential gene expression in APL cells undergoing differentiation after ATRA treatment (27, 28). However, total RNA sequencing captures entire transcriptomes and can thus provide a more thorough exploration of how ATRA differentially affects the gene expression of ATRA-responsive and ATRA-resistance cells. Here, we performed RNA-Seq on APL cells treated with both ATRA and TSA to identify transcriptional changes potentially associated with mechanical response to treatment. As a control, we performed the same treatments and RNA-Seq analysis on HL-60 cells, which were derived from a patient diagnosed with APL but whose cells lacked the PML-RARA fusion gene (56). We performed RNA-Seq on these three cell lines (AP-1060, NB4, HL-60) under different conditions (untreated, TSA, ATRA, TSA + ATRA), and identified differentially expressed genes between samples and conditions (57).

In comparing cell lines, ATRA-sensitive NB4 had the most differentially expressed genes (2,523) between its untreated and ATRA-treated samples (Supplementary Fig. S9A). AP-1060 had fewer differentially expressed genes (1,907), reflecting its ATRA resistance, and HL-60 had the least (1,176), reflecting its lack of sensitivity to ATRA due to a lack of the PML-RARA fusion gene (Supplementary Fig. S9A). In contrast, there was only one differentially expressed gene (C1QA, part of the classical complement pathway) between TSA-treated and untreated NB4 compared to 86 in TSA-treated vs. untreated AP-1060 (Supplementary Fig. S9B). We also evaluated the interaction of TSA with ATRA by comparing gene expression of ATRA-treated cells to cells treated with both TSA and ATRA. This analysis revealed 13 differentially expressed genes in NB4, compared to 385 in AP-1060 (Supplementary Fig. S9D, S9E). The greater number of differentially expressed genes in AP-1060 compared to NB4, both when evaluating TSA treatment and when combining TSA and ATRA, underscores the cell line’s sensitivity to TSA HDAC inhibition that we previously measured with mechano-NPS.

We performed principal component analysis on all differentially expressed genes and evaluated the loadings of individual genes to each principal component (Fig. 5A, Supplementary Fig. S10). The first two principal components (PC1 and PC2) clearly separate the different cell types. The third principal component (PC3) appears to specifically separate only AP-1060 samples treated with TSA. Analyzing the top 100 highest-loading genes for each principal component, PC3 was especially enriched for gene ontology terms associated with immune cell function and activation compared to PC1 and PC2. PC1 was primarily enriched for cellular reorganization and RNA processing. PC2 was enriched for several biological processes including metabolism and cell activation (Supplementary Fig. S10).

**Figure 5:**
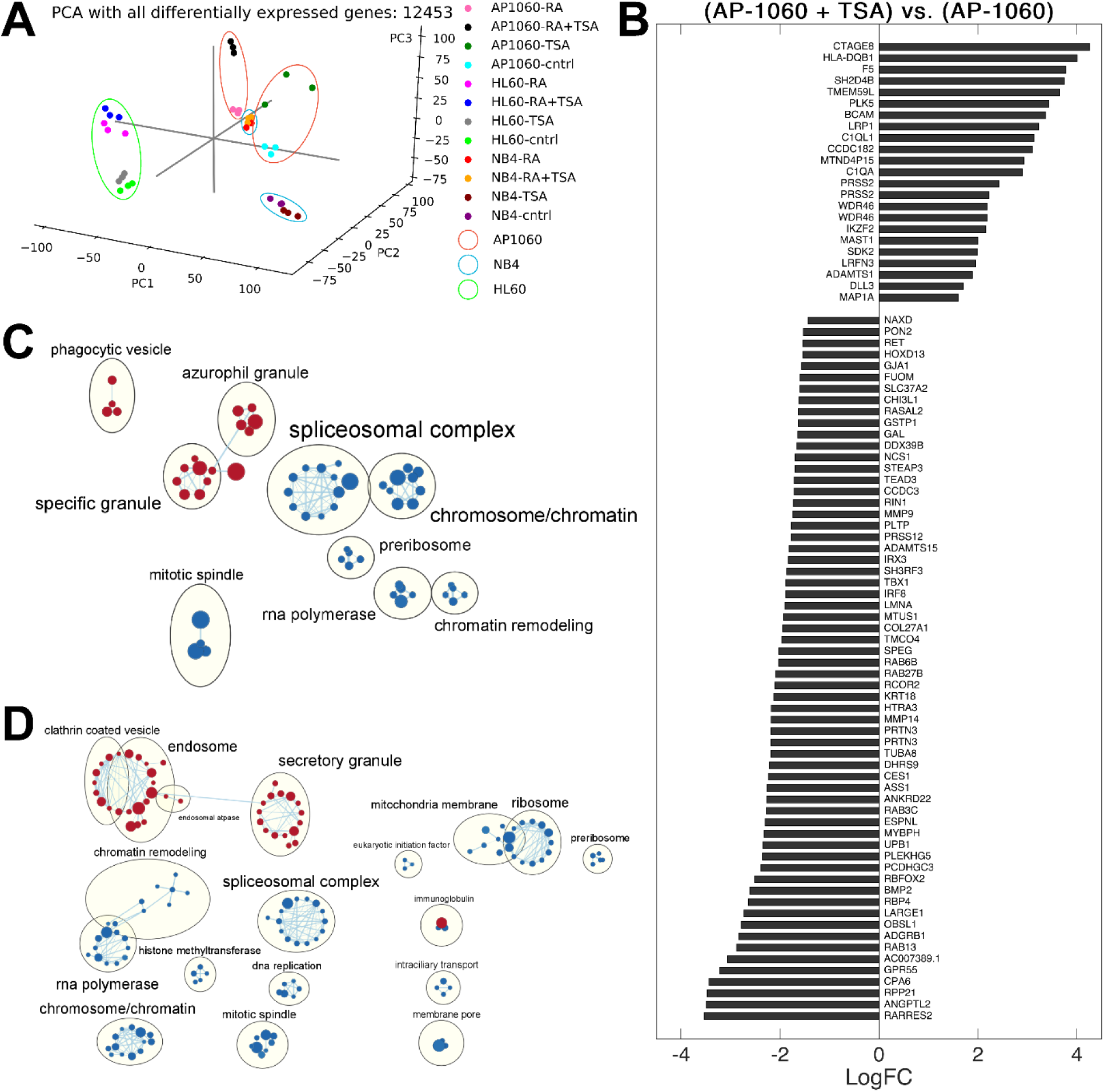
RNAseq and differential expression analysis of APL cells highlights transcriptional differences in ATRA-resistant AP-1060 cells. **A**. Principal component analysis of counts per differentially expressed gene identified in any of 66 between-sample pairwise comparisons. **B**. Differentially expressed genes identified in comparing TSA-treated vs. untreated AP-1060. **C**. Enrichment map of positively (red) and negatively (blue) enriched gene sets for ATRA-treated AP-1060 cells. GSEA was performed from the *GO_CC* (Gene ontology cellular component) collection. **D**. Enrichment map of positively (red) and negatively (blue) enriched gene sets for ATRA-treated NB4 cells. GSEA was performed from the *GO_CC* (Gene ontology cellular component) collection.

We analyzed the expression of the 86 differentially expressed genes between untreated AP-1060 and TSA-treated AP-1060. The retinoid signaling-related genes RARRES2 and RBP4 were among the most highly downregulated genes (1^st^ and 11^th^, respectively) (Fig. 5B). RARRES2 was found to be upregulated in an HL-60 population made multi-drug resistant via selection with ATRA, and RBP4 has been shown to promote the extracellular transport of retinoic acid (58, 59). As such, downregulation of these two genes could increase sensitivity to ATRA. We filtered these 86 genes by association to the cytoskeleton to identify genes that might directly contribute to cell structure and, by extension, mechanical properties. Several microtubule-associated genes were downregulated, as were OBSL1, MYBPH, EPSNL, ANKRD22, KRT18, and LMNA (in decreasing order of log-fold change in expression). Only microtubule-associated MAP1A was upregulated. Overall, we would expect that the downregulation of cytoskeletal genes contributes to the softening of TSA-treated AP-1060 cells.

We next investigated the differentially expressed genes in AP-1060 and NB4 after 96 hours of ATRA treatment. First, we compared our differential expression results to cDNA microarray results from Yang *et al*. where NB4 cells were treated with ATRA for 96 hours, finding agreement in 24 of 62 upregulated genes and 15 of 43 downregulated genes (28). The remaining genes found to be differentially expressed by Yang *et al*. were not found to be differentially expressed in our comparisons. Gene set enrichment analysis (GSEA) comparing ATRA-treated and untreated AP-1060 and NB4 cells reflected the more dramatic transcriptional response in NB4 (Fig. 5C, 5D; Supplementary Fig. S9A). Positively enriched gene sets in ATRA-treated AP-1060 involved granulocytic functions (phagocytosis, granules), and negatively enriched gene sets linked to transcription, RNA processing, and chromosome/chromatin organization (Fig. 5C). These same gene sets were similarly enriched in ATRA-treated NB4. Additionally, negatively enriched gene sets in ATRA-treated NB4 involved components of DNA replication and epigenetic regulation, including histone methyltransferases (Fig. 5D). As we showed that DNA content (Fig. 3) and chromatin decondensation (Fig. 4) both modulate mechanical phenotype, transcriptional changes affecting chromatin organization and DNA replication may be important drivers of cell softening in ATRA-treated NB4.

Last, we constructed a gene set to specifically investigate the immunophenotypic landscape of differentiating AP-1060 and NB4 cells. As expected, NB4 cells exhibited greater upregulation of innate immunity-related genes (indicative of differentiation) after ATRA treatment compared to AP-1060 (Supplementary Fig. S11). These data provide a more thorough transcriptomic analysis of ATRA-induced differentiation compared to prior work using cDNA microarrays (27, 28), as well as how this response differs in an ATRA-resistant cell line.

## Discussion and Conclusions

APL is a highly aggressive acute myeloid leukemia subtype that can quickly lead to patient death if therapeutic ATRA regimens are unsuccessful. ATRA induces differentiation of malignant immature promyelocytes, which will otherwise proliferate to the point of patient morbidity. Using mechano-NPS, we have, for the first time, uncovered specific cell mechanical properties that correlate with ATRA-resistant APL. Specifically, we determined that ATRA-resistant AP-1060 cells are stiffer and more viscous than ATRA-sensitive NB4 cells. Through an in-depth study, we determined that the mechanics of APL cells are more dependent on the mechanical properties of their nuclei rather than the F-actin network. Specifically, major changes in DNA content strongly influenced cell mechanical phenotypes, with greater amounts of nuclear DNA being associated with stiffer and more viscous cells. Chromatin decondensation had a subtler effect on APL cells but still led to the softening of ATRA-resistant AP-1060 cells, a response that was observed in ATRA-treated NB4 but not AP-1060 cells. Through RNA-Seq, we discovered enriched gene sets that may also affect mechanical phenotype by way of chromatin remodeling.

The importance of nuclear mechanics in APL is made more evident when considering our results with cells that were synchronized into S-phase and had their F-actin networks disrupted. When we inhibited HDAC activity with TSA to induce chromatin remodeling, we found that chromatin decondensation can also significantly affect nuclear mechanics. Synchronized and actin-destabilized AP-1060 cells treated with TSA were more deformable and less viscous than similarly treated NB4 cells. ATRA treatment stiffened these AP-1060 cells, in contrast to cells that were solely treated with ATRA. This may be a consequence of AP-1060’s ATRA resistance and the unique epigenomic state of S-phase synchronized cells, where decondensed chromatin is more accessible and susceptible to retinoid signaling (60). However, further work is warranted to identify the exact mechanism behind this unexpected synergistic effect between cell cycle synchronization and ATRA. Unlike AP-1060 cells, synchronized NB4 cells with disrupted F-actin networks consistently softened after ATRA treatment. Overall, we conclude that while NB4 is robustly sensitive to ATRA, AP-1060 is uniquely sensitive to HDAC inhibition with TSA.

Identifying the biophysical factors that link a particular pathological phenotype to a specific mechanical phenotype is a new approach toward the implementation of physical and mechanical biomarkers to characterizing cancer cells. Understanding these factors can help explain the biological underpinnings of the mechanical properties of cancer cells, whose potential clinical applications are frequently discussed (61, 62). To this point, we systematically perturbed subcellular components of APL cells and assessed their relevance to the mechanical phenotype associated with ATRA resistance. In our RNA-Seq analysis, we highlighted genes and gene sets associated with our observations of APL cells and their mechanical properties, and also provide a resource characterizing the transcriptome of differentiating and ATRA-resistant APL cells. We bridge together two critically under-examined aspects of APL, ultimately uncovering a previously unknown relationship between drug resistance and mechanical phenotype.

## Materials and Methods

### Device design and fabrication

All mechano-NPS devices utilized in these studies consisted of 12.9 ± 0.1 µm high microfluidic channel molded into a polydimethylsiloxane (PDMS) slab that was bonded to a glass substrate with pre-defined platinum (Pt) electrodes and gold (Au) contact pads. PDMS slabs included in-line filters with a pore size of 20 µm to prevent the ingress of cell clusters that would clog the channel. The central contraction segment, 7 µm × 2000 µm (W × L), was flanked by a single node-segment at its entrance and 10 recovery segments at its exit. While all nodes were 85 µm × 50 µm (W × L), the single segment located in front of the contraction segment was 13 µm × 800 µm (W × L) and the recovery segments were each 13 µm × 290 µm (W × L). Contraction segment length was chosen such that cells experienced deformation for ∼150-200 ms. The series of 10 recovery segments provided up to ∼500 ms of sampling for recovering cells, which was sufficient to observe the exponential decay of cell strain to a steady-state value.

Microfluidic channels were fabricated using a standard soft lithography process, and Pt/Au electrodes were fabricated using a lift-off process with electron-gun evaporation for metal deposition. A detailed fabrication process for mechano-NPS devices is provided in Supplementary Note 2.

### Mechano-node pore sensing

Mechano-node pore sensing was performed as previously described (5). A DC potential (< 3 V) is applied across the mechano-NPS channel and a four-terminal measurement is performed. Cells transiting the channel partially block the flow of electric current, leading to a modulated current pulse. In the experiments performed, cell suspensions were prepared at a concentration of 300,000 cells/mL in 1X phosphate buffered saline (PBS) solution supplemented with 2% fetal bovine serum (FBS, VWR 89510-186) to reduce cell-cell and cell-PDMS adhesion. Cell suspensions were injected into mechanoNPS devices using a microfluidic pressure controller (Elveflow OB1) with a nominal inlet pressure of 80 mbar. Signals were sampled at 50 kHz, post-processed with a moving-average low pass filter, and then downsampled to 2.5 kHz. A custom command-line interface program, written in MATLAB <u>(available on GitHub)</u> rapidly identifies mechanoNPS pulses, determines subpulse features (i.e. magnitude and duration), and extracts physical and mechanical parameters.

### Cell culture

AP-1060 cells, obtained from Dr. S. Kogan, University of California-San Francisco, San Franscisco, CA, U.S.A., were cultured in 70% IMDM (Gibco 12440053) supplemented with 20% FBS, 1% Penicillin-Streptomycin (Gibco 15070063), and 10% conditioned medium from cell line 5637 (ATCC HTB-9). For conditioned medium from cell line 5637, cells were seeded in 10 mL of RPMI-1640 (Corning 10-040-CV) supplemented with 10% FBS and 1% Penicillin-Streptomycin (Gibco 15070063). Medium was exchanged after 24 hours, then collected at 48 hours after initial seeding. Conditioned medium was then sterilized using a 0.22 µm polyethylsufone filter (Millipore Sigma SLGPM33RS). NB4 (DSMZ ACC 207) and HL-60 (ATCC CCL-240) cells were cultured in 90% RPMI-1640 supplemented with 10% FBS and 1% Penicillin-Streptomycin. AP-1060 cells were passaged at a density of 2 × 10^6^ cells/mL, seeded in wells of a 24-well plate at a density of 1 × 10^6^, and maintained at 37 °C in 5% CO_2_. NB4 and HL-60 cells were passaged at a density of 5 × 10^5^ cells/mL, seeded in wells of a 24-well plate at a density of 1 × 10^6^, and maintained at 37 °C in 5% CO_2_.

### Pharmacological treatments

Dry powder all-trans retinoic acid (ATRA, Sigma-Aldrich R2625) was reconstituted in anhydrous dimethylsulfoxide to 25 mg/mL for an 83.3 mM concentrated stock solution (Sigma-Aldrich 276855). Concentrated stock solution was aliquoted and snap-frozen in liquid nitrogen, then transferred to a −80 °C freezer for long-term storage. Concentrated stock ATRA was thawed and diluted to 10 mM in dimethyl sulfoxide (DMSO) prior to use, then further diluted to working concentration (10 nM to 10 µM for testing ATRA response; 1 µM otherwise) in cell-culture media. LatrunculinA (LatA, Abcam ab144290) was reconstituted in ethyl alcohol (Sigma-Aldrich 459844) and added to cell-culture media at a concentration of 0.5 µg/mL for 30 minutes to disrupt actin filaments. 100 nM of Trichostatin A (TSA, Fisher Scientific 14-061, Mfg. Tocris Bioscience), reconstituted in DMSO, was added to cells for 24 hours to inhibit histone deacetylase activity. For synchronized and LatA-treated cells, cell cultures were first synchronized in S-phase (see below). Cells experiencing no further treatment were treated with LatA (see above) immediately prior to mechano-NPS analysis. Cells treated with TSA were incubated in cell culture media supplemented with 100 nM TSA for 24 hours, then treated with LatA (see above) immediately prior to mechano-NPS analysis. Cells treated with ATRA were incubated in cell culture media supplemented with 1 µM ATRA at 0 and 48 hours after the completion of cell-cycle synchronization, then treated with LatA (see above) 96 hours after synchronization, followed by immediate analysis with mechano-NPS. For all pharmacological treatments, cells were pelleted via centrifugation at 200 rcf for 5 minutes, rinsed once with 1X PBS, then pelleted again, and resuspended for mechano-NPS analysis (see above).

### Cell-cycle arrest

A double thymidine block was used to arrest cells at S-phase. Cells were resuspended at 1 × 10^6^ cells/mL in cell-culture media supplemented with 2 mM thymidine (Abcam ab143719) and incubated at 37 °C in 5% CO_2_ for 18 hours. They were then isolated via centrifugation at 200 rcf for 5 minutes, resuspended at the same density in cell-culture media supplemented with 10 µM deoxycytidine (Abcam ab146218), and subsequently incubated at 37 °C in 5% CO_2_ for 8 hours. Finally, cells were again collected via centrifugation at 200 rcf for 5 minutes, resuspended at the same density in cell-culture media supplemented with 2 mM thymidine, and incubated at 37 °C in 5% CO_2_ for 18 hours. Synchronized cells were then isolated via centrifugation at 200 rcf for 5 minutes and rinsed once with 1X PBS for further experiments. Colcemid solution in 1X PBS (Gibco 15212012) was used to arrest cell division and synchronize cells in M-phase. Colcemid was added directly to cells in culture media at a concentration of 1 µg/mL. Cell cultures were then incubated at 37 °C in 5% CO_2_ for 2 hours. Synchronized cells were then isolated via centrifugation at 200 rcf for 5 minutes and rinsed once with 1X PBS for further experiments. Cell cycle arrest in either S-or M-phase was confirmed via flow cytometric analysis of DNA content (see below).

### Fluorescence staining and immunostaining

Silicone isolators (Grace BioLabs CWS-13R-0.5) were pressed onto poly-L-lysine glass slides (VWR 16002-116) to form small wells. Cells in suspension culture were resuspended at 5 × 10^5^ cells/mL in 1X PBS, then pipetted into the wells, and allowed to settle and adhere for 1 hour while incubating at 37 °C in 5% CO_2_. Wells were then washed with 1X PBS to remove unbound cells. Cells were then fixed with 4% paraformaldehyde (Sigma-Aldrich P6148) and permeabilized with 0.1% Triton X-100 (Sigma-Aldrich T8787). For immunocytochemistry, cells were blocked with donkey serum (Sigma-Aldrich D9663) for 1 hour at room temperature, washed with 1X PBS, then stained overnight with anti-LaminA primary antibody (Invitrogen MA1-06101) at a ratio of 1:200 at 4 °C. Fluorescence staining solutions for immunocytochemistry contained 40 µM Hoechst 33342 (Thermo Scientific 62249), 165 nM rhodamine phalloidin (Biotium 00027), and 1:1000 donkey anti-mouse Alexa-Fluor 488-conjugated secondary antibody (Invitrogen A-21202). Fluorescence staining solutions for ATRA and LatA experiments contained the same concentrations of Hoechst 33342 and rhodamine phalloidin.

### Flow cytometric analysis

Cells were isolated by centrifugation at 200 rcf for 5 minutes and resuspended in 1X PBS. For assessing DNA content, cells were stained with 40 µM Hoechst 33342 for 30 minutes, washed with 1X PBS, and analyzed on a BD LSR Fortessa X20 with BD FACSDiva 9.0 software. For immunostained flow cytometry, cells were blocked with an anti-Fc Receptor polyclonal antibody (Invitrogen 14-9161-73) for 30 minutes. An antibody mix for a three-color flow cytometry panel was prepared, consisting of FITC-anti-CD11b (Biolegend 101206), Super Bright 600-anti-CD18 (Invitrogen 63-0189-41), and PerCP-eFluor 710-anti-CD52 (Invitrogen 46-0529-41). Samples were incubated with fluorophore-conjugated antibodies for 35 minutes, then washed several times with 1X PBS. Samples were then stained with LIVE/DEAD Fixable Violet (Invitrogen L34955) and washed several times with 1X PBS prior to analysis. For each experimental condition, one unstained control and three fluorescence-minus-one controls for each antibody-conjugated fluorophore were prepared (Supplementary Fig. S2). Single-stain positive control samples for each fluorophore were prepared prior to all analysis to compute a compensation matrix. Compensation was set such that the median fluorescence intensity in negative channels was the same across single-stain control samples. All analysis of flow cytometry data was performed using Cytobank Community (63).

### RNA-Seq and differential expression analysis

RNA was extracted from ∼1 M cells using the QIAGEN RNeasy Mini Kit (QIAGEN 74104). A total of 36 samples were prepared: for each of three cell lines (AP-1060, HL-60, and NB4), the four conditions of untreated, ATRA, TSA, and ATRA & TSA were run in triplicate. Libraries were prepared using NEBNext Ultra II RNA Library Prep Kit for Illumina (New England Biolabs E7770S) and the NEBNext Poly(A) mRNA Magnetic Isolation Module (New England Biolabs E7490S) with 1 *μ*g total RNA input. Paired-end 2 × 100 bp sequencing was performed on three Illumina HiSeq 4000 lanes with twelve samples per lane for an average sequencing depth of 31M reads per sample. Transcripts were quantified using Salmon (version 0.10.0) (64) in mapping-based mode with the human reference transcriptome (Ensembl GRCh38 version 86) (65). Read counts were then converted to counts per gene using Bioconductor (version 3.7) (66) with the *tximport* package (67). Differential expression was determined using *edgeR* (version 3.22.0) and *limma* (version 3.32.0) (68–72). Significance in differential expression was then filtered by a log-fold change greater than 1 using the TREAT method (*treat* function from *limma*) (57). Principal component analysis was performed with the *scikit-learn* library in Python (73), and gene set overlaps were computed using the Molecular Signatures Database (MSigDB) and its *Investigate Gene Sets* tools (74, 75). Gene set enrichment analysis was carried out using GSEA 4.0.3, using 497 gene sets from GO Cellular Component (c5.go.cc.v7.2.symbols.gmt) (74, 76) after applying a maximum and minimum size threshold of 500 and 15 genes, respectively. For enrichment mapping, enriched gene set nodes were filtered using a false discovery rate cutoff of 25%, and edges were filtered using a similarity cutoff of 50% (77, 78).

### High-speed imaging

100 mbar of nominal inlet pressure from an Elveflow OB1 pressure controller drove cells through a mechanoNPS device. Images were acquired using a Fastec IL-5 high-speed camera at 1500 frames per second with a 167 µs shutter speed. Movie playback was set to 50 frames per second.

### Mechano-NPS statistical analysis

Statistical tests for mechano-NPS data were applied to measurements of both *wCDI* and recovery time. Significant differences were principally calculated using two-sample Student’s t-tests with a significance criterion of *α* = 0.05. For experiments involving multiple comparisons, test statistics were instead computed using a Tukey’s range test.

### Power analysis

To quantify the power of all mechano-NPS statistical tests, we performed a *post hoc* power analysis on all statistical tests made for both *wCDI* and recovery time (see Supplementary Tables 1 and 2). Statistical power was calculated according to a two-sample Student’s t-test. Where applicable, a Bonferroni correction to the significance criterion (by default, α = 0.05) was made for experiments involving multiple comparisons. The statistical power for each test given the measured effect size and sample size was calculated and reported. For tests not exceeding a power value of 0.80, we reported the minimum effect size needed for a statistical power of 0.80, as well as the measured effect size.

## Acknowledgements

This work was supported by the National Institutes of Health (NIH 1R01CA190843-01) and B. Li and A. Maslan were both supported by National Science Foundation Graduate Research Fellowships. We also thank S. Kogan for generously providing the AP-1060 cell line; J. Kim, M. Kozminsky, and A. Dong for their insights and discussions; R. Rex, M. Kozminsky, and H. Randolph for their critical reading of the manuscript; and the UC Berkeley Flow Cytometry Facility (Berkeley, CA), which provided access to flow cytometers and analysis software.

## References

1. Gossett, D.R., H.T.K. Tse, S.A. Lee, Y. Ying, A.G. Lindgren, O.O. Yang, J. Rao, A.T. Clark, and D. Di Carlo. 2012. Hydrodynamic stretching of single cells for large population mechanical phenotyping. Proc. Natl. Acad. Sci. U. S. A. 109:7630–7635.

2. Tse, H.T.K., D.R. Gossett, Y.S. Moon, M. Masaeli, M. Sohsman, Y. Ying, K. Mislick, R.P. Adams, J. Rao, and D. Di Carlo. 2013. Quantitative diagnosis of malignant pleural effusions by single-cell mechanophenotyping. Sci. Transl. Med. 5.

3. Byun, S., S. Son, D. Amodei, N. Cermak, J. Shaw, J.H. Kang, V.C. Hecht, M.M. Winslow, T. Jacks, P. Mallick, and S.R. Manalis. 2013. Characterizing deformability and surface friction of cancer cells. Proc. Natl. Acad. Sci. U. S. A. 110:7580–7585.

4. Toepfner, N., C. Herold, O. Otto, P. Rosendahl, J. Sta, L. Menschner, A. Jacobi, M. Kra, B. Henriques-normark, N. Tregay, P. Mellroth, E.R. Chilvers, R. Berner, M. Suttorp, and M. Bornha. 2018. Detection of human disease conditions by phenotyping of blood. 1–22.

5. Kim, J., S. Han, A. Lei, M. Miyano, J. Bloom, V. Srivastava, M.R. Stampfer, Z.J. Gartner, M.A. LaBarge, and L.L. Sohn. 2018. Characterizing cellular mechanical phenotypes with mechano-node-pore sensing. Microsystems Nanoeng. 4:1–12.

6. Kimmerling, R.J., S.M. Prakadan, A.J. Gupta, N.L. Calistri, M.M. Stevens, S. Olcum, N. Cermak, R.S. Drake, K. Pelton, F. De Smet, K.L. Ligon, A.K. Shalek, and S.R. Manalis. 2018. Linking single-cell measurements of mass, growth rate, and gene expression. Genome Biol. 19:207.

7. Lam, W.A., M.J. Rosenbluth, and D.A. Fletcher. 2007. Chemotherapy exposure increases leukemia cell stiffness. Blood. 109:3505–3508.

8. Rosenbluth, M.J., W.A. Lam, and D.A. Fletcher. 2006. Force microscopy of nonadherent cells: A comparison of leukemia cell deformability. Biophys. J. 90:2994–3003.

9. Lautenschläger, F., S. Paschke, S. Schinkinger, A. Bruel, M. Beil, and J. Guck. 2009. The regulatory role of cell mechanics for migration of differentiating myeloid cells. Proc. Natl. Acad. Sci. U. S. A. 106:15696–15701.

10. Stein, E., B. McMahon, H. Kwaan, J.K. Altman, O. Frankfurt, and M.S. Tallman. 2009. The coagulopathy of acute promyelocytic leukaemia revisited. Best Pract. Res. Clin. Haematol. 22:153–163.

11. Pals, S.T., D.J.J. de Gorter, and M. Spaargaren. 2007. Lymphoma dissemination: the other face of lymphocyte homing. Blood. 110:3102–3111.

12. Stevens, M.M., C.L. Maire, N. Chou, M.A. Murakami, D.S. Knoff, Y. Kikuchi, R.J. Kimmerling, H. Liu, S. Haidar, N.L. Calistri, N. Cermak, S. Olcum, N.A. Cordero, A. Idbaih, P.Y. Wen, D.M. Weinstock, K.L. Ligon, and S.R. Manalis. 2016. Drug sensitivity of single cancer cells is predicted by changes in mass accumulation rate. Nat. Biotechnol. 34:1161–1167.

13. Mu, L., J.H. Kang, S. Olcum, K.R. Payer, N.L. Calistri, R.J. Kimmerling, S.R. Manalis, and T.P. Miettinen. 2020. Mass measurements during lymphocytic leukemia cell polyploidization decouple cell cycle-and cell size-dependent growth. Proc. Natl. Acad. Sci. 117:15659–15665.

14. Kang, J.H., T.P. Miettinen, L. Chen, S. Olcum, G. Katsikis, P.S. Doyle, and S.R. Manalis. 2019. Noninvasive monitoring of single-cell mechanics by acoustic scattering. Nat. Methods. 16:263–269.

15. Stevens, M.M., C.L. Maire, N. Chou, M.A. Murakami, D.S. Knoff, Y. Kikuchi, R.J. Kimmerling, H. Liu, S. Haidar, N.L. Calistri, N. Cermak, S. Olcum, N.A. Cordero, A. Idbaih, P.Y. Wen, D.M. Weinstock, K.L. Ligon, and S.R. Manalis. 2016. Drug sensitivity of single cancer cells is predicted by changes in mass accumulation rate. Nat. Biotechnol. 34:1161–1167.

16. Rowley, J., H. Golomb, and C. Dougherty. 1977. 15/17 TRANSLOCATION, A CONSISTENT CHROMOSOMAL CHANGE IN ACUTE PROMYELOCYTIC LEUKAEMIA. Lancet. 309:549–550.

17. Puccetti, E., and M. Ruthardt. 2004. Acute promyelocytic leukemia: PML/RARα and the leukemic stem cell. Leukemia. 18:1169–1175.

18. Mendez, L., M. Chen, and P.P. Pandolfi. 2018. Molecular Genetics of APL. In: Acute Promyelocytic Leukemia. Cham: Springer International Publishing. pp. 41–53.

19. Tallman, M.S., J.W. Andersen, C.A. Schiffer, F.R. Appelbaum, J.H. Feusner, A. Ogden, L. Shepherd, C. Willman, C.D. Bloomfield, J.M. Rowe, and P.H. Wiernik. 1997. All-trans -Retinoic Acid in Acute Promyelocytic Leukemia. N. Engl. J. Med. 337:1021–1028.

20. Douer, D., L.N. Zickl, C.A. Schiffer, F.R. Appelbaum, J.H. Feusner, L. Shepherd, C.L. Willman, C.D. Bloomfield, E. Paietta, R.E. Gallagher, J.H. Park, J.M. Rowe, P.H. Wiernik, and M.S. Tallman. 2013. All-trans retinoic acid and late relapses in acute promyelocytic leukemia: Very long-term follow-up of the North American Intergroup Study I0129. Leuk. Res. 37:795–801.

21. Park, J.H., B. Qiao, K.S. Panageas, M.J. Schymura, J.G. Jurcic, T.L. Rosenblat, J.K. Altman, D. Douer, J.M. Rowe, and M.S. Tallman. 2011. Early death rate in acute promyelocytic leukemia remains high despite all-trans retinoic acid. Blood. 118:1248– 1254.

22. Ribeiro, R.C., and E. Rego. 2006. Management of APL in developing countries: epidemiology, challenges and opportunities for international collaboration. Hematology Am. Soc. Hematol. Educ. Program. 162–168.

23. Douer, D., S. Preston-Martin, E. Chang, P.W. Nichols, K.J. Watkins, and A.M. Levine. 1996. High frequency of acute promyelocytic leukemia among Latinos with acute myeloid leukemia. Blood. 87:308–313.

24. Stein, E., B. McMahon, H. Kwaan, J.K. Altman, O. Frankfurt, and M.S. Tallman. 2009. The coagulopathy of acute promyelocytic leukaemia revisited. Best Pract. Res. Clin. Haematol. 22:153–163.

25. Gallagher, R. 2002. Retinoic acid resistance in acute promyelocytic leukemia. Leukemia. 16:1940–1958.

26. Noguera, N.I., G. Catalano, C. Banella, M. Divona, I. Faraoni, T. Ottone, W. Arcese, and M.T. Voso. 2019. Acute Promyelocytic Leukemia: Update on the Mechanisms of Leukemogenesis, Resistance and on Innovative Treatment Strategies. Cancers (Basel). 11.

27. Lee, K.H., M.Y. Chang, J.I. Ahn, D.H. Yu, S.S. Jung, J.H. Choi, Y.H. Noh, Y.S. Lee, and M.J. Ahn. 2002. Differential gene expression in retinoic acid-induced differentiation of acute promyelocytic leukemia cells, NB4 and HL-60 cells. Biochem. Biophys. Res. Commun. 296:1125–1133.

28. Yang, L., H. Zhao, S.-W. Li, K. Ahrens, C. Collins, S. Eckenrode, Q. Ruan, R.A. McIndoe, and J.-X. She. 2003. Gene Expression Profiling during All-trans Retinoic Acid-Induced Cell Differentiation of Acute Promyelocytic Leukemia Cells. J. Mol. Diagnostics. 5:212–221.

29. Roussel, M.J.S., and M. Lanotte. 2001. Maturation sensitive and resistant t(15;17) NB4 cell lines as tools for APL physiopathology: Nomenclature of cells and repertory of their known genetic alterations and phenotypes. Oncogene. 20:7287–7291.

30. Lanotte, M., V. Martin-Thouvenin, S. Najman, P. Balerini, F. Valensi, and R. Berger. 1991. NB4, a maturation inducible cell line with t(15;17) marker isolated from a human acute promyelocytic leukemia (M3). Blood. 77:1080–1086.

31. Sun, Y., S.H. Kim, D.C. Zhou, W. Ding, E. Paietta, F. Guidez, A. Zelent, K.H. Ramesh, L. Cannizzaro, R.P. Warrell, and R.E. Gallagher. 2004. Acute promyelocytic leukemia cell line AP-1060 established as a cytokine-dependent culture from a patient clinically resistant to all-trans retinoic acid and arsenic trioxide. Leukemia. 18:1258–1269.

32. May, R.C., and L.M. Machesky. 2001. Phagocytosis and the actin cytoskeleton. J. Cell Sci. 114:1061–1077.

33. Dahl, K.N., A.J.S. Ribeiro, and J. Lammerding. 2008. Nuclear Shape, Mechanics, and Mechanotransduction. Circ. Res. 102:1307–1318.

34. Friedl, P., K. Wolf, and J. Lammerding. 2011. Nuclear mechanics during cell migration. Curr. Opin. Cell Biol. 23:55–64.

35. Wolf, K., M. te Lindert, M. Krause, S. Alexander, J. te Riet, A.L. Willis, R.M. Hoffman, C.G. Figdor, S.J. Weiss, and P. Friedl. 2013. Physical limits of cell migration: Control by ECM space and nuclear deformation and tuning by proteolysis and traction force. J. Cell Biol. 201:1069–1084.

36. Lawrence, S.M., R. Corriden, and V. Nizet. 2018. The Ontogeny of a Neutrophil: Mechanisms of Granulopoiesis and Homeostasis. Microbiol. Mol. Biol. Rev. 82.

37. Rowat, A.C., D.E. Jaalouk, M. Zwerger, W.L. Ung, I.A. Eydelnant, D.E. Olins, A.L. Olins, H. Herrmann, D.A. Weitz, and J. Lammerding. 2013. Nuclear Envelope Composition Determines the Ability of Neutrophil-type Cells to Passage through Micron-scale Constrictions. J. Biol. Chem. 288:8610–8618.

38. Lammerding, J., P.C. Schulze, T. Takahashi, S. Kozlov, T. Sullivan, R.D. Kamm, C.L. Stewart, and R.T. Lee. 2004. Lamin A/C deficiency causes defective nuclear mechanics and mechanotransduction. J. Clin. Invest. 113:370–378.

39. Yamaguchi, H., and J. Condeelis. 2007. Regulation of the actin cytoskeleton in cancer cell migration and invasion. Biochim. Biophys. Acta - Mol. Cell Res. 1773:642–652.

40. Pollard, T.D., and J.A. Cooper. 2009. Actin, a Central Player in Cell Shape and Movement. Science (80-.). 326:1208–1212.

41. Chalut, K.J., and E.K. Paluch. 2016. The Actin Cortex: A Bridge between Cell Shape and Function. Dev. Cell. 38:571–573.

42. Gardel, M.L., I.C. Schneider, Y. Aratyn-Schaus, and C.M. Waterman. 2010. Mechanical Integration of Actin and Adhesion Dynamics in Cell Migration. Annu. Rev. Cell Dev. Biol. 26:315–333.

43. Pegoraro, A.F., P. Janmey, and D.A. Weitz. 2017. Mechanical Properties of the Cytoskeleton and Cells. Cold Spring Harb. Perspect. Biol. 9:a022038.

44. Gabriele, S., A.-M. Benoliel, P. Bongrand, and O. Théodoly. 2009. Microfluidic Investigation Reveals Distinct Roles for Actin Cytoskeleton and Myosin II Activity in Capillary Leukocyte Trafficking. Biophys. J. 96:4308–4318.

45. Webster, M., K.L. Witkin, and O. Cohen-Fix. 2009. Sizing up the nucleus: nuclear shape, size and nuclear-envelope assembly. J. Cell Sci. 122:1477–1486.

46. Aureille, J., V. Buffière-Ribot, B.E. Harvey, C. Boyault, L. Pernet, T. Andersen, G. Bacola, M. Balland, S. Fraboulet, L. Van Landeghem, and C. Guilluy. 2019. Nuclear envelope deformation controls cell cycle progression in response to mechanical force. EMBO Rep. 20.

47. Chu, F.-Y., S.C. Haley, and A. Zidovska. 2017. On the origin of shape fluctuations of the cell nucleus. Proc. Natl. Acad. Sci. 114:10338–10343.

48. Otto, O., P. Rosendahl, A. Mietke, S. Golfier, C. Herold, D. Klaue, S. Girardo, S. Pagliara, A. Ekpenyong, A. Jacobi, M. Wobus, N. Töpfner, U.F. Keyser, J. Mansfeld, E. Fischer-Friedrich, and J. Guck. 2015. Real-time deformability cytometry: On-the-fly cell mechanical phenotyping. Nat. Methods. 12:199–202.

49. Bostock, C. 1971. An evaluation of the double thymidine block for synchronizing mammalian cells at the G1-S border*1. Exp. Cell Res. 68:163–168.

50. Bernardi, R., and P.P. Pandolfi. 2007. Structure, dynamics and functions of promyelocytic leukaemia nuclear bodies. Nat. Rev. Mol. Cell Biol. 8:1006–1016.

51. Nervi, C., F.F. Ferrara, M. Fanelli, M.R. Rippo, B. Tomassini, P.F. Ferrucci, M. Ruthardt, V. Gelmetti, C. Gambacorti-Passerini, D. Diverio, F. Grignani, P.G. Pelicci, and R. Testi. 1998. Caspases mediate retinoic acid-induced degradation of the acute promyelocytic leukemia PML/RARα fusion protein. Blood. 92:2244–2251.

52. Stephens, A.D., E.J. Banigan, S.A. Adam, R.D. Goldman, and J.F. Marko. 2017. Chromatin and lamin a determine two different mechanical response regimes of the cell nucleus. Mol. Biol. Cell. 28:1984–1996.

53. Stephens, A.D., P.Z. Liu, E.J. Banigan, L.M. Almassalha, V. Backman, S.A. Adam, R.D. Goldman, and J.F. Marko. 2018. Chromatin histone modifications and rigidity affect nuclear morphology independent of lamins. Mol. Biol. Cell. 29:220–233.

54. Minucci, S., C. Nervi, F. Lo Coco, and P.G. Pelicci. 2001. Histone deacetylases: a common molecular target for differentiation treatment of acute myeloid leukemias? Oncogene. 20:3110–3115.

55. Ceccacci, E., and S. Minucci. 2016. Inhibition of histone deacetylases in cancer therapy: Lessons from leukaemia. Br. J. Cancer. 114:605–611.

56. Dalton, W.J., M. Ahearn, K. McCredie, E. Freireich, S. Stass, and J. Trujillo. 1988. HL-60 cell line was derived from a patient with FAB-M2 and not FAB-M3. Blood. 71:242–247.

57. Mccarthy, D.J., and G.K. Smyth. 2009. Testing significance relative to a fold-change threshold is a TREAT. Bioinformatics. 25:765–771.

58. Liu, S.-M., W. Chen, and J. Wang. 2015. Distinguishing between cancer cell differentiation and resistance induced by all-trans retinoic acid using transcriptional profiles and functional pathway analysis. Sci. Rep. 4:5577.

59. Blaner, W.S. 2019. Vitamin A signaling and homeostasis in obesity, diabetes, and metabolic disorders. Pharmacol. Ther. 197:153–178.

60. Alabert, C., and A. Groth. 2012. Chromatin replication and epigenome maintenance. Nat. Rev. Mol. Cell Biol. 13:153–167.

61. Kozminsky, M., and L.L. Sohn. 2020. The promise of single-cell mechanophenotyping for clinical applications. Biomicrofluidics. 14:031301.

62. Carey, T.R., K.L. Cotner, B. Li, and L.L. Sohn. 2019. Developments in label-free microfluidic methods for single-cell analysis and sorting. Wiley Interdiscip. Rev. Nanomedicine Nanobiotechnology. 11:e1529.

63. Kotecha, N., P.O. Krutzik, and J.M. Irish. 2010. Web-Based Analysis and Publication of Flow Cytometry Experiments. Curr. Protoc. Cytom. 53:10.17.1-10.17.24.

64. Patro, R., G. Duggal, M.I. Love, R.A. Irizarry, C. Kingsford, and C. Biology. 2017. Salmon. 14:417–419.

65. Cunningham, F., P. Achuthan, W. Akanni, J. Allen, M.R. Amode, I.M. Armean, R. Bennett, J. Bhai, K. Billis, S. Boddu, C. Cummins, C. Davidson, K.J. Dodiya, A. Gall, C. G. Girón, L. Gil, T. Grego, L. Haggerty, E. Haskell, T. Hourlier, O.G. Izuogu, S.H. Janacek, T. Juettemann, M. Kay, M.R. Laird, I. Lavidas, Z. Liu, J.E. Loveland, J.C. Marugán, T. Maurel, A.C. McMahon, B. Moore, J. Morales, J.M. Mudge, M. Nuhn, D. Ogeh, A. Parker, A. Parton, M. Patricio, A.I. Abdul Salam, B.M. Schmitt, H. Schuilenburg, D. Sheppard, H. Sparrow, E. Stapleton, M. Szuba, K. Taylor, G. Threadgold, A. Thormann, A. Vullo, B. Walts, A. Winterbottom, A. Zadissa, M. Chakiachvili, A. Frankish, S.E. Hunt, M. Kostadima, N. Langridge, F.J. Martin, M. Muffato, E. Perry, M. Ruffier, D.M. Staines, S.J. Trevanion, B.L. Aken, A.D. Yates, D.R. Zerbino, and P. Flicek. 2019. Ensembl 2019. Nucleic Acids Res. 47:D745–D751.

66. Huber, W., V.J. Carey, R. Gentleman, S. Anders, M. Carlson, B.S. Carvalho, H.C. Bravo, S. Davis, L. Gatto, T. Girke, R. Gottardo, F. Hahne, K.D. Hansen, R.A. Irizarry, M. Lawrence, M.I. Love, J. MacDonald, V. Obenchain, A.K. Oleś, H. Pagès, A. Reyes, P. Shannon, G.K. Smyth, D. Tenenbaum, L. Waldron, and M. Morgan. 2015. Orchestrating high-throughput genomic analysis with Bioconductor. Nat. Methods. 12:115–121.

67. Soneson, C., M.I. Love, and M.D. Robinson. 2015. Differential analyses for RNA-seq: transcript-level estimates improve gene-level inferences. F1000Research. 4:1521.

68. Robinson, M.D., D.J. McCarthy, and G.K. Smyth. 2010. edgeR: a Bioconductor package for differential expression analysis of digital gene expression data. Bioinformatics. 26:139–40.

69. McCarthy, D.J., Y. Chen, and G.K. Smyth. 2012. Differential expression analysis of multifactor RNA-Seq experiments with respect to biological variation. Nucleic Acids Res. 40:4288–97.

70. Phipson, B., S. Lee, I.J. Majewski, W.S. Alexander, and G. Smyth. 2016. Robust Hyperparameter Estimation Protects. Ann. Appl. Stat. 10:946–963.

71. Ritchie, M.E., B. Phipson, D. Wu, Y. Hu, C.W. Law, W. Shi, and G.K. Smyth. 2015. Limma powers differential expression analyses for RNA-sequencing and microarray studies. Nucleic Acids Res. 43:e47.

72. Law, C.W., Y. Chen, W. Shi, and G.K. Smyth. 2014. voom: precision weights unlock linear model analysis tools for RNA-seq read counts. Genome Biol. 15:R29.

73. Pedregosa, F., G. Varoquaux, A. Gramfort, V. Michel, B. Thirion, O. Grisel, M. Blondel, A. Müller, J. Nothman, G. Louppe, P. Prettenhofer, R. Weiss, V. Dubourg, J. Vanderplas, A. Passos, D. Cournapeau, M. Brucher, M. Perrot, and É. Duchesnay. 2012. Scikit-learn: Machine Learning in Python..

74. Subramanian, A., P. Tamayo, V.K. Mootha, S. Mukherjee, B.L. Ebert, M.A. Gillette, A. Paulovich, S.L. Pomeroy, T.R. Golub, E.S. Lander, and J.P. Mesirov. 2005. Gene set enrichment analysis: A knowledge-based approach for interpreting genome-wide expression profiles. Proc. Natl. Acad. Sci. 102:15545–15550.

75. Liberzon, A., A. Subramanian, R. Pinchback, H. Thorvaldsdottir, P. Tamayo, and J.P. Mesirov. 2011. Molecular signatures database (MSigDB) 3.0. Bioinformatics. 27:1739– 1740.

76. Mootha, V.K., C.M. Lindgren, K.-F. Eriksson, A. Subramanian, S. Sihag, J. Lehar, P. Puigserver, E. Carlsson, M. Ridderstråle, E. Laurila, N. Houstis, M.J. Daly, N. Patterson, J.P. Mesirov, T.R. Golub, P. Tamayo, B. Spiegelman, E.S. Lander, J.N. Hirschhorn, D. Altshuler, and L.C. Groop. 2003. PGC-1α-responsive genes involved in oxidative phosphorylation are coordinately downregulated in human diabetes. Nat. Genet. 34:267– 273.

77. Merico, D., R. Isserlin, O. Stueker, A. Emili, and G.D. Bader. 2010. Enrichment Map: A Network-Based Method for Gene-Set Enrichment Visualization and Interpretation. PLoS One. 5:e13984.

78. Shannon, P. 2003. Cytoscape: A Software Environment for Integrated Models of Biomolecular Interaction Networks. Genome Res. 13:2498–2504.

