## Supplementary Information for "Mechanical phenotyping of acute promyelocytic leukemia reveals unique biomechanical responses in retinoic acid-resistant populations"

**Supplementary Note 1: Quantifying rates of cell recovery from deformation**

The cell can be approximated as a homogeneous sphere of a Kelvin-Voigt material. Its release from the constriction channel into recovery segments constitutes a step change in applied compressive stress. Consequently, the cell strain will relax to a steady-state value according to the following exponential decay model:

$\varepsilon\left( t \right)=\varepsilon_{0}\exp(-t/\tau)+\varepsilon_{\infty}$	( 1 )

where $\varepsilon(t)$ is the strain over time after the step change in stress, $\varepsilon_{0}$ is the strain at the time of the step change in stress, $\tau$ is the time constant of the exponential decay, and $\varepsilon_{\infty}$ is the steady-state value for strain after the cell has fully relaxed. In node-pore sensing techniques, the magnitude of the current drop is related to the size of the particle [1,2]. A transform of variables from the stress-strain space to the voltage-current space gives the following equation governing the measurement of this viscoelastic creep in mechano-NPS:

$\Delta I\left( t \right)=\Delta I_{0}\left[ 1-\exp\left( -t/\tau\right) \right]+{\Delta I}_{\infty}$	( 2 )

where $\Delta I$ is equivalent to a difference in current from the baseline: $I_{baseline}-I(t)$, $\Delta I_{0}$ is the current drop associated with a cell at the time of release from the constriction channel, and $\Delta I_{\infty}$ is the current drop associated with a cell at its steady-state strain after complete relaxation. Rewriting Eq. 2 as a function of measured current rather than current drop yields:

$I\left( t \right)=I_{0} exp(-t/\tau)+I_{\infty}$	( 3 )

where $I(t)$ is the measured current over time after the step change in stress, $I_{0}$ is the current associated with the cell at the time of release from the constriction channel, $\tau$ is the time constant of the exponential decay in voltage, and $I_{\infty}$ is the current associated with the cell at its steady-state strain after complete relaxation.

Mechano-NPS subpulses from after the cell exits the constriction channel provide values for $I(t)$ and $t$ to which we fit a linear model to estimate $\tau$. To perform linear least squares regression for this model, we linearize Eq. 3 by subtracting $I_{\infty}$ from $I\left( t \right)$ and taking the logarithm of both sides:

$\ln\left[ I\left( t \right)-I_{\infty} \right]=\ln I_{0}-t/\tau$	( 4 )

After performing linear least squares, we can then use the slope of the linear model to calculate a confidence interval for the time constant of the exponential decay, thus estimating a cell’s rate of recovery from deformation.

**Supplementary Note 2: Detailed fabrication process flow for mechano-NPS devices**

A standard soft lithography process was used to create the PDMS microfluidic channels. Briefly, negative relief structures were fabricated onto polished silicon wafer using SU8-3010 epoxy resist (MicroChem). SU-8 3010 was spun at 1850 rpm for 30 seconds and baked at 95 $^{\circ}$C for 8 minutes. The resist-coated wafer was then exposed to a mask with UV light at a dose of 160 mJ/cm^2^, baked again at 65 $^{\circ}$C for 1 minute, and then at 95 $^{\circ}$C for 3 minutes. Finally, the wafer was immersed in SU-8 developer (MicroChem) for 2 minutes, then rinsed with water and dried. This process yielded a film thickness and microchannel height of 12.9 ± 0.1 µm (mean ± std. dev.). Sylgard 184 PDMS (Dow Corning) pre-polymer and curing agent were mixed in a ratio of 9:1, degassed, and then poured onto the negative-relief masters. After curing for 2 hours at 85 $^{\circ}$C, PDMS slabs with the embedded microfluidic channels were cut and peeled from the relief masters, cored with a 1.5 mm biopsy punch (Harris Uni-Core, Fisher Scientific) to provide input/output access to the microchannel, and then cleaned with isopropanol and deionized water (DI, 18 MΩ).

The Pt electrodes and Au contact pads were patterned onto glass slides using standard photolithography. First, glass slides were patterned with positive-tone S1813 resist (MicroChem) by spin-coating at 3000 rpm and soft-baking at 100 $^{\circ}$C for 1 minute. The wafer was then exposed to a mask with UV light at a dose of 300 mJ/cm^2^, and subsequently developed in MF-321 developer (MicroChem) for 45 seconds. A thin metal film consisting of 75 Å Ti, 250 Å Pt, and 250 Å Au was then deposited onto the patterned glass slides using electron-gun evaporation. Excess metal and photoresist were lifted off using acetone., The fabricated metal electrodes were then cleaned with acetone, isopropanol, and DI water. The glass slides with pre-fabricated electrodes and the molded PDMS devices were simultaneously treated with oxygen plasma (2 minutes, 450 mTorr, 30 W, Harrick Plasma) before being bonded together and baked at 125 $^{\circ}$C for 5 minutes.

**Supplementary Table 1: Power analysis for wCDI by mechano-NPS experimental groups**

| Group 1 | Group 2 | N_1_ | N_2_ | Power | Min. effect size | Actual effect size |
| --- | --- | --- | --- | --- | --- | --- |
| AP-1060 | NB4 | 246 | 124 | 1.00 | N/A | N/A |
| AP-1060 | + ATRA | 246 | 333 | 0.08 | 0.033 | 0.012 |
| NB4 | + ATRA | 124 | 402 | 1.00 | N/A | N/A |
| AP-1060 | + Colcemid | 614 | 236 | 1.00 | N/A | N/A |
| NB4 | + Colcemid | 123 | 167 | 1.00 | N/A | N/A |
| AP-1060 Sync + LatA | + TSA | 155 | 84 | 1.00 | N/A | N/A |
| AP-1060 Sync + LatA | + ATRA | 155 | 199 | 1.00 | N/A | N/A |
| AP-1060 Sync + Lat A + TSA | + ATRA | 84 | 199 | 1.00 | N/A | N/A |
| NB4 Sync  + LatA | + TSA | 75 | 114 | 0.68 | 0.094 | 0.083 |
| NB4 Sync  + LatA | + ATRA | 75 | 95 | 0.79 | 0.103 | 0.101 |
| NB4 Sync  + LatA + TSA | + ATRA | 114 | 95 | 0.09 | 0.058 | 0.019 |
| AP-1060 | + LatA | 268 | 327 | 0.05 | 0.023 | 0.002 |
| NB4 | + LatA | 341 | 239 | 0.13 | 0.055 | 0.016 |

*Post-hoc* power analysis was performed on all statistical tests for mechano-NPS measurements of *wCDI*. Table rows correspond to specific comparisons made between experimental Group 1 and Group 2 with respective sample sizes N_1_ and N_2_. Bonferroni corrections to the default significance criterion $\alpha$ = 0.05 were made for experiments with multiple comparisons. For tests with statistical power equal to or less than 0.80, a minimum effect size for $\pi$ = 0.80 and actual effect size are reported.

**Supplementary Table 2: Power analysis for recovery time by mechano-NPS experimental groups**

| Group 1 | Group 2 | N_1_ | N_2_ | Power | Min. effect size | Actual effect size |
| --- | --- | --- | --- | --- | --- | --- |
| AP-1060 | NB4 | 246 | 124 | 1.00 | N/A | N/A |
| AP-1060 | + ATRA | 246 | 333 | 0.96 | N/A | N/A |
| NB4 | + ATRA | 124 | 402 | 0.01 | 27.7 | 1.09 |
| AP-1060 | + Colcemid | 614 | 236 | 0.13 | 33.6 | 9.64 |
| NB4 | + Colcemid | 123 | 167 | 1.00 | N/A | N/A |
| AP-1060 Sync + LatA | + TSA | 155 | 84 | 0.27 | 81.4 | 45.1 |
| AP-1060 Sync + LatA | + ATRA | 155 | 199 | 0.29 | 52.6 | 29.9 |
| AP-1060 Sync + Lat A + TSA | + ATRA | 84 | 199 | 0.11 | 42.2 | 15.2 |
| NB4 Sync + LatA | + TSA | 75 | 114 | 0.44 | 55.9 | 38.9 |
| NB4 Sync + LatA | + ATRA | 75 | 95 | 0.47 | 61.3 | 43.8 |
| NB4 Sync  + LatA + TSA | + ATRA | 114 | 95 | 1.00 | N/A | N/A |
| AP-1060 | + LatA | 268 | 327 | 1.00 | N/A | N/A |
| NB4 | + LatA | 341 | 239 | 0.80 | 38.2 | 38.0 |

*Post-hoc* power analysis was performed on all statistical tests for mechano-NPS measurements of recovery time constant. Table rows correspond to specific comparisons made between experimental Group 1 and Group 2 with respective sample sizes N_1_ and N_2_. Bonferroni corrections to the default significance criterion $\alpha$ = 0.05 were made for experiments with multiple comparisons. For tests with statistical power equal to or less than 0.80, a minimum effect size for $\pi$ = 0.80 and actual effect size are reported.

**Supplementary Movie 1:** AP-1060 cell transiting the section of a mechano-NPS channel used for initial measurements of cell size and unconstrained velocity. Segment (narrow channel) width is 13 µm, and the node (wide channel) width is 85 µm.

**Supplementary Movie 2:** AP-1060 entering the contraction segment of a mechano-NPS channel where cells are deformed in transit. Segment width is 7 µm, and the node (wide channel) width is 85 µm.

**Supplementary Movie 3:** AP-1060 exiting the constriction channel of a mechano-NPS channel. Cells exit the constriction channel deformed into a bullet shape due to a combination of shear stress and compressive stresses from the constriction channel side walls. Mechano-NPS measures the relaxation from this bullet shape to a more spherical morphology to quantify a cell’s recovery from deformation. Contraction segment width is 7 microns, node (wide channel) width is 85 µm, and recovery segment width is 13 µm.

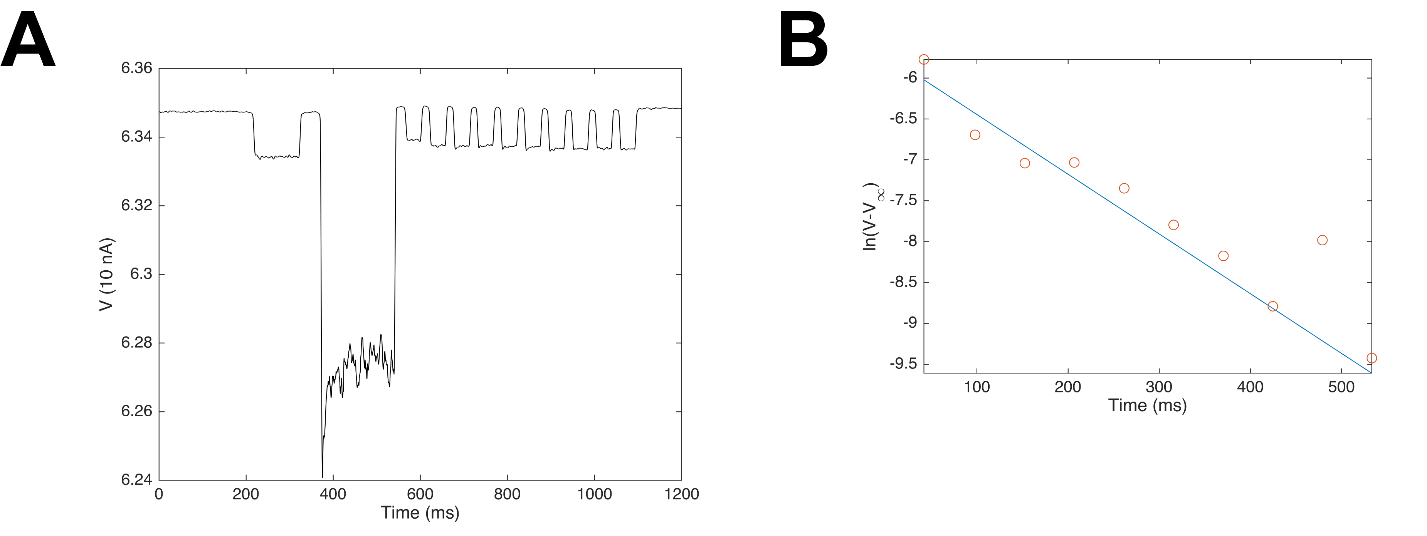
**Supplementary Figure S1:** MATLAB processing of mechano-NPS signals (**A**) for quantifying cell recovery from deformation. **A.** Pulses are pre-processed as previously published [Kim *et al* 2018]., using a low-pass filter, base-line normalization, and derivative threshold to identify the start and end of each subpulse **B**. The logarithm of voltage values for each recovery segment subpulse (red circles) are fit to a linear function (blue line) using linear least squares regression (see Supplementary Note 2).

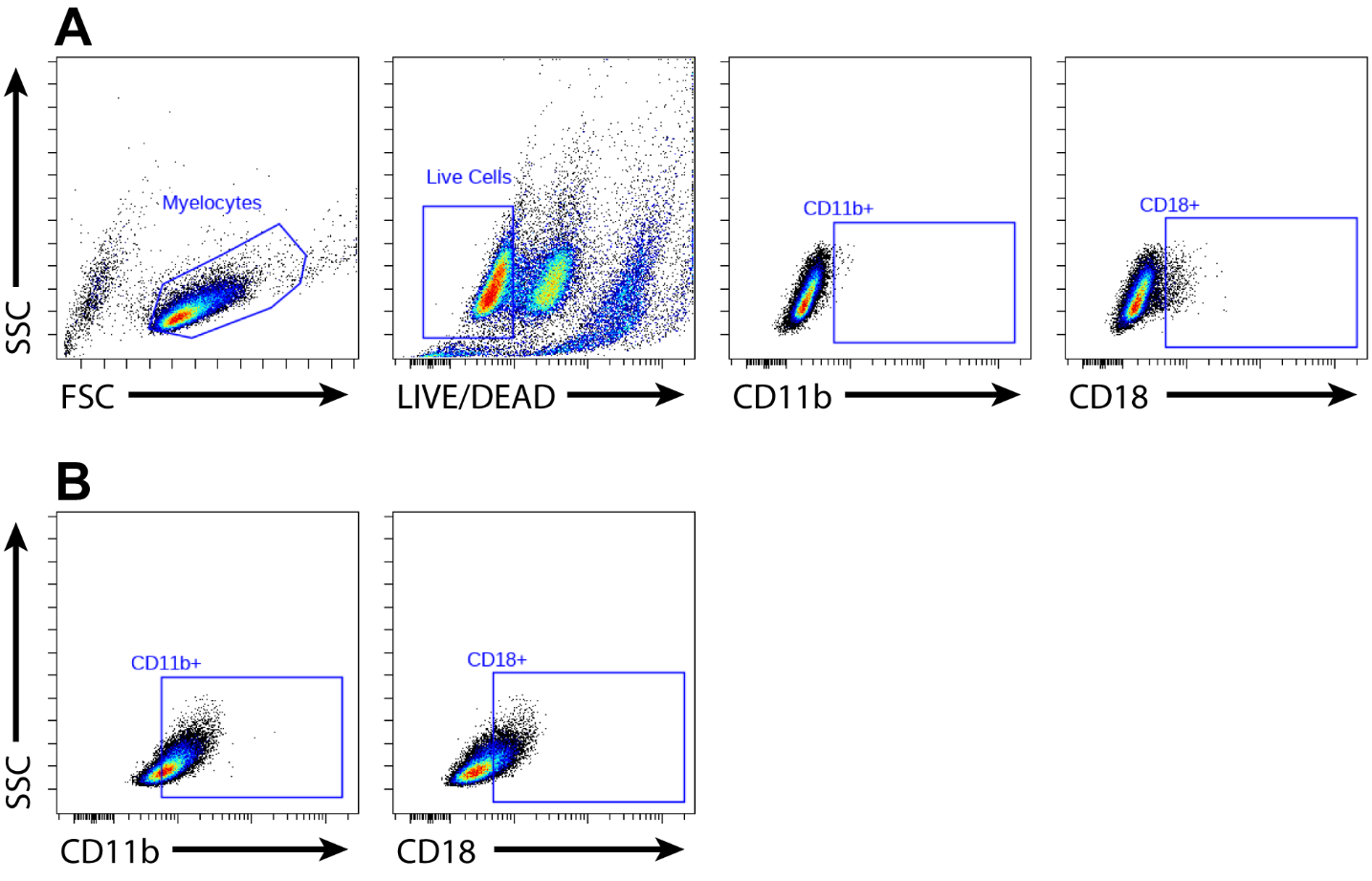
**Supplementary Figure S2:** Gating strategy for flow cytometry measurements of the CD11b and CD18 expression of AP-1060 and NB4 cells induced to differentiate with ATRA. **A.** Myelocytes are first identified as a cluster in side scatter (SSC) v. forward scatter (FSC). Live cells are then gated by the LIVE/DEAD-low cluster based on fluorescence from the LIVE/DEAD Violet dye. Fluorescence-minus-one (FMO) controls for each marker were used to determine the true negative fluorescence intensity distribution for the markers’ respective fluorophores. **B.** Using gates established by FMO controls, the proportion of ATRA-differentiated cells expressing a certain marker can be measured.

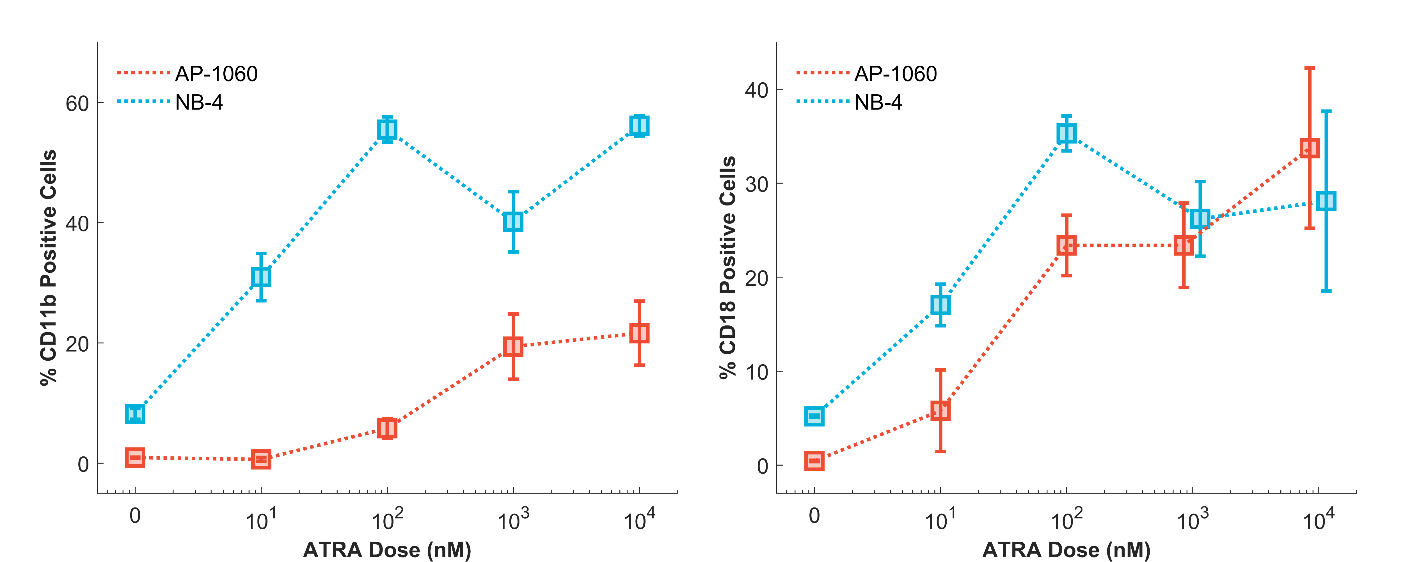

**Supplementary Figure S3:** Proportions of cells positive for CD11b and CD18, markers associated with mature/differentiated myelocytes, after treatment with varying doses of ATRA. A stronger differentiation response is seen in NB4 cells, indicating a higher susceptibility to induced differentiation via ATRA.

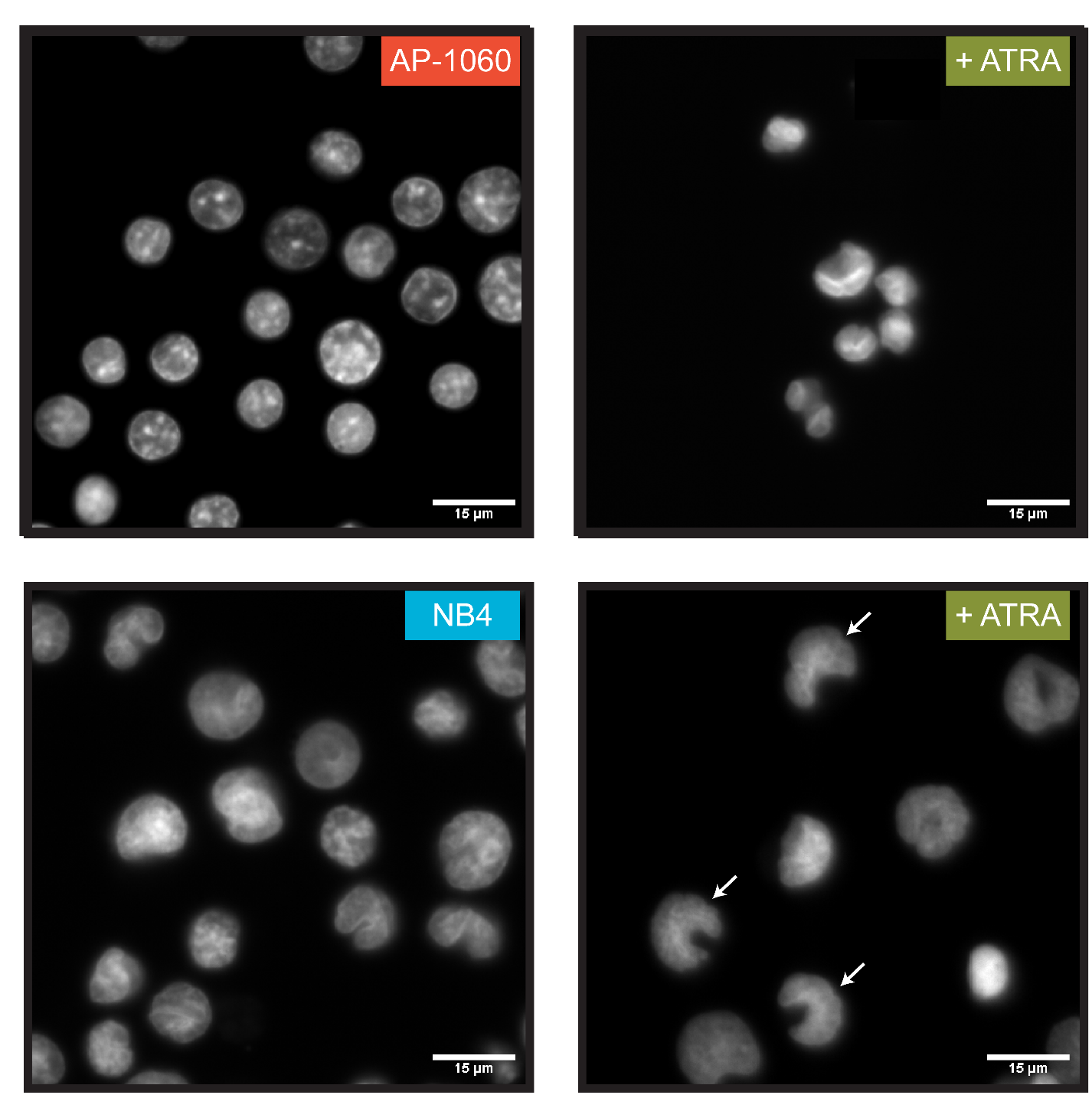
**Supplementary Figure S4:** Fluorescence microscopy of AP-1060 and NB4 nuclei before and after ATRA treatment. Nuclei were stained with Hoechst 33342 dye without counterstain. White arrows represent lobulated nuclei.

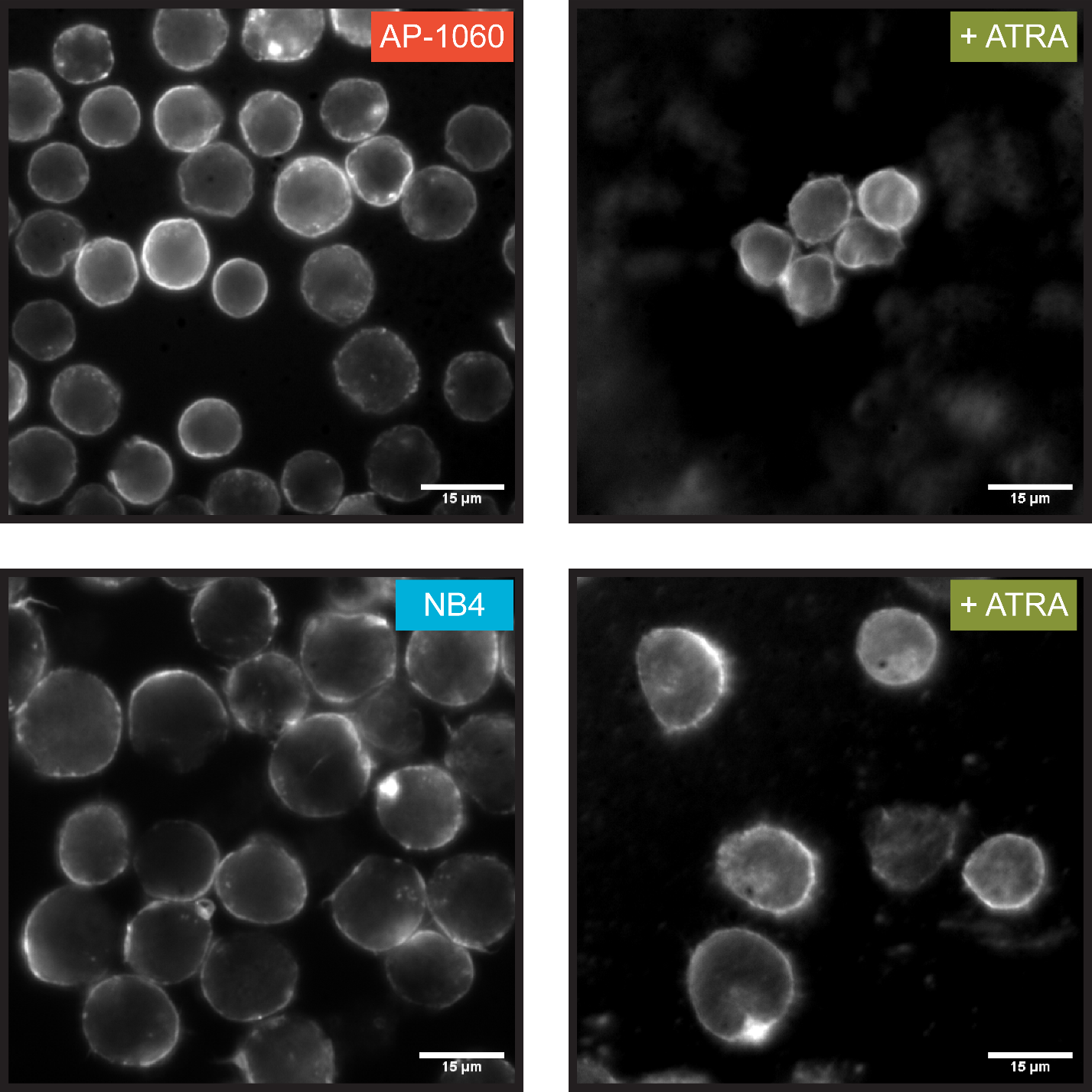

**Supplementary Figure S5:** Isolation of rhodamine phalloidin channel from Figure 2C.

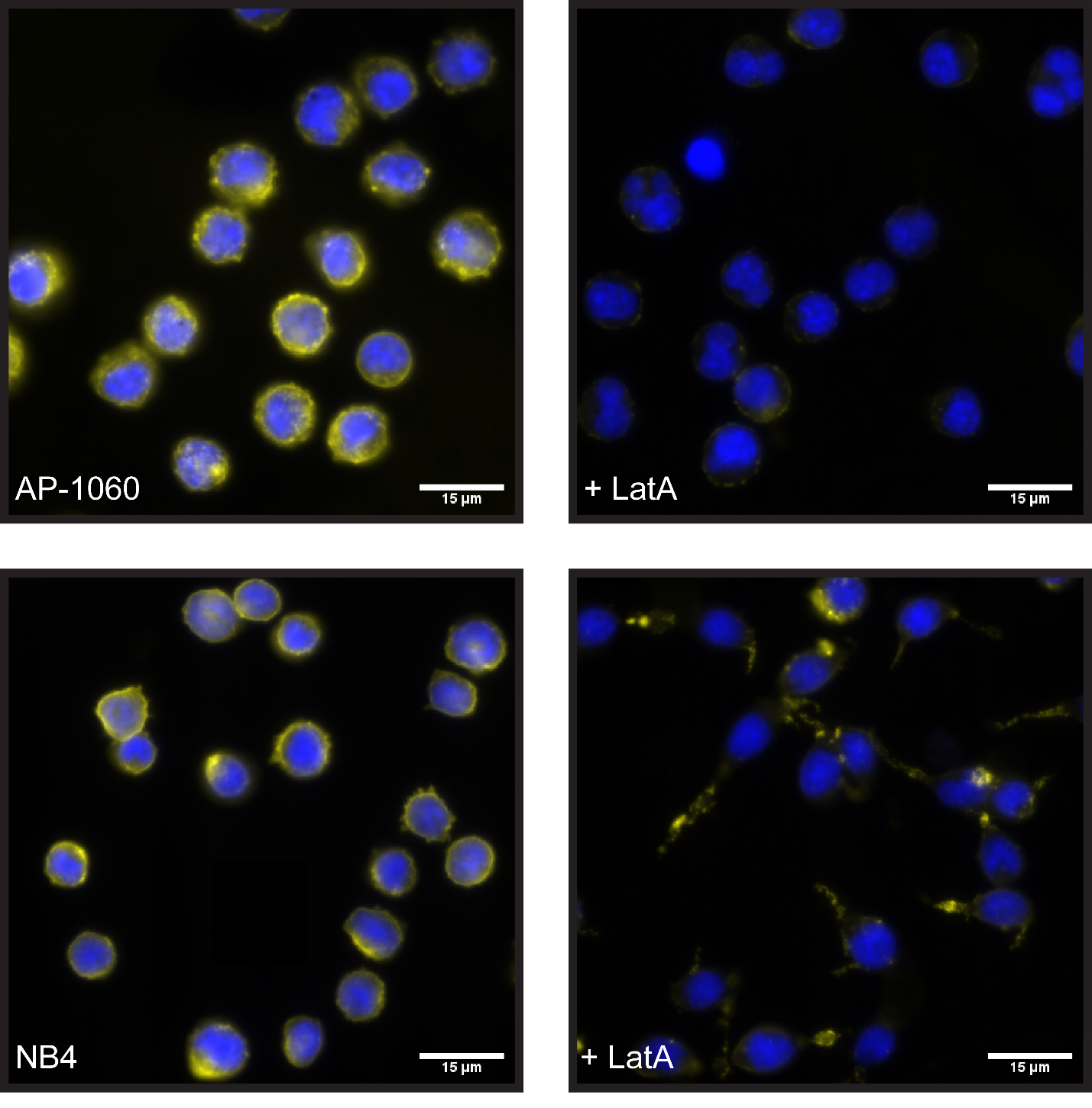
**Supplementary Figure S6:** Fluorescence microscopy of AP-1060 and NB4 cells before and after treatment with Latrunculin A. Blue represents DNA stained with Hoechst 33342 dye, and yellow represents actin stained with rhodamine phalloidin.

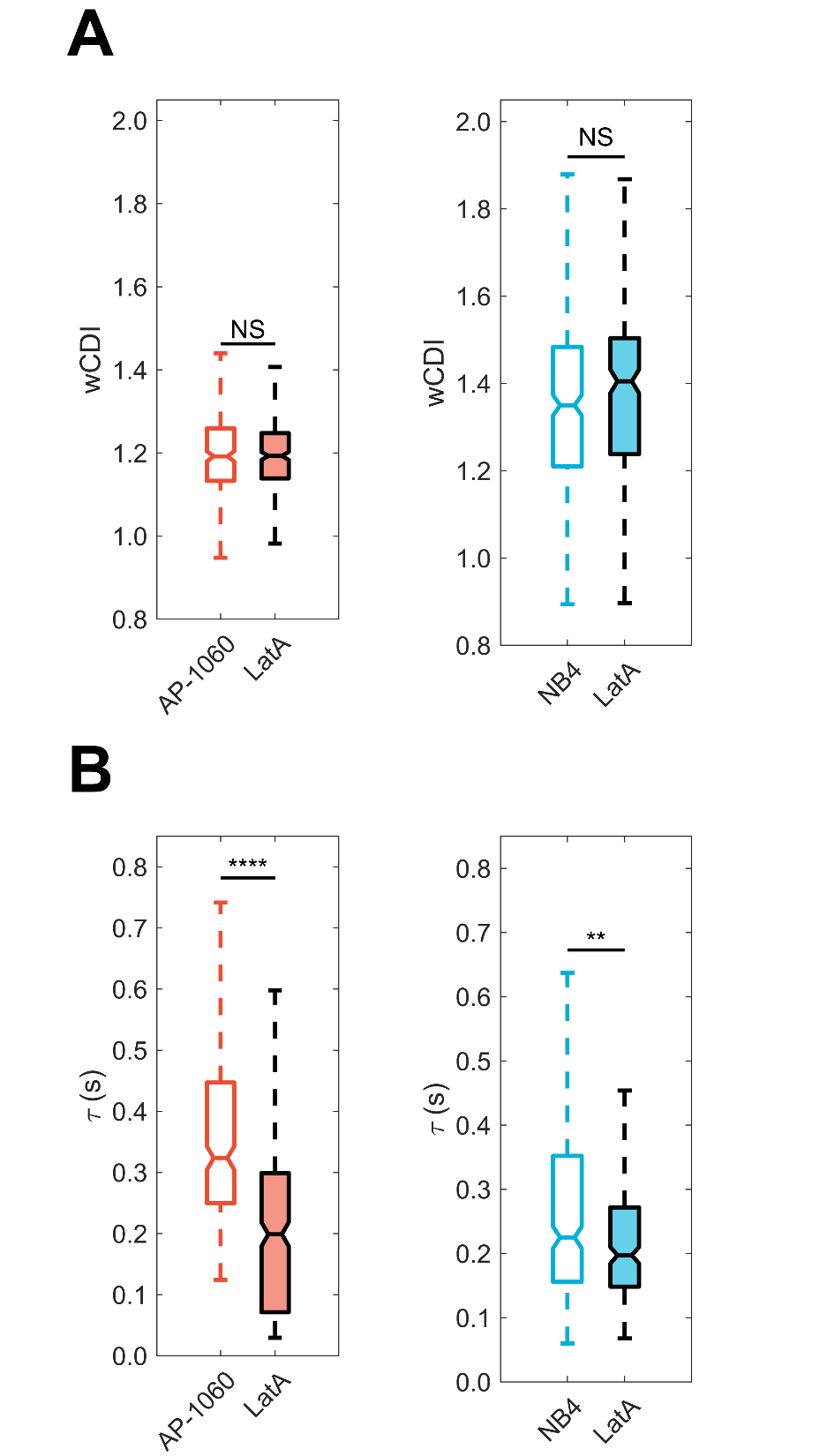

**Supplementary Figure S7: A, B.** Box plots of wCDI (**A**) and recovery times (**B**) for AP-1060 and NB4 cells before and after treatment with latrunculin A. Notches represent 95% confidence intervals for the true median of each distribution. *n* $\geq$ 239 for each distribution across 3 different devices. ***p* < 0.01 *****p* < 0.0001, NS no significance; determined by two-sample Student’s *t*-tests.

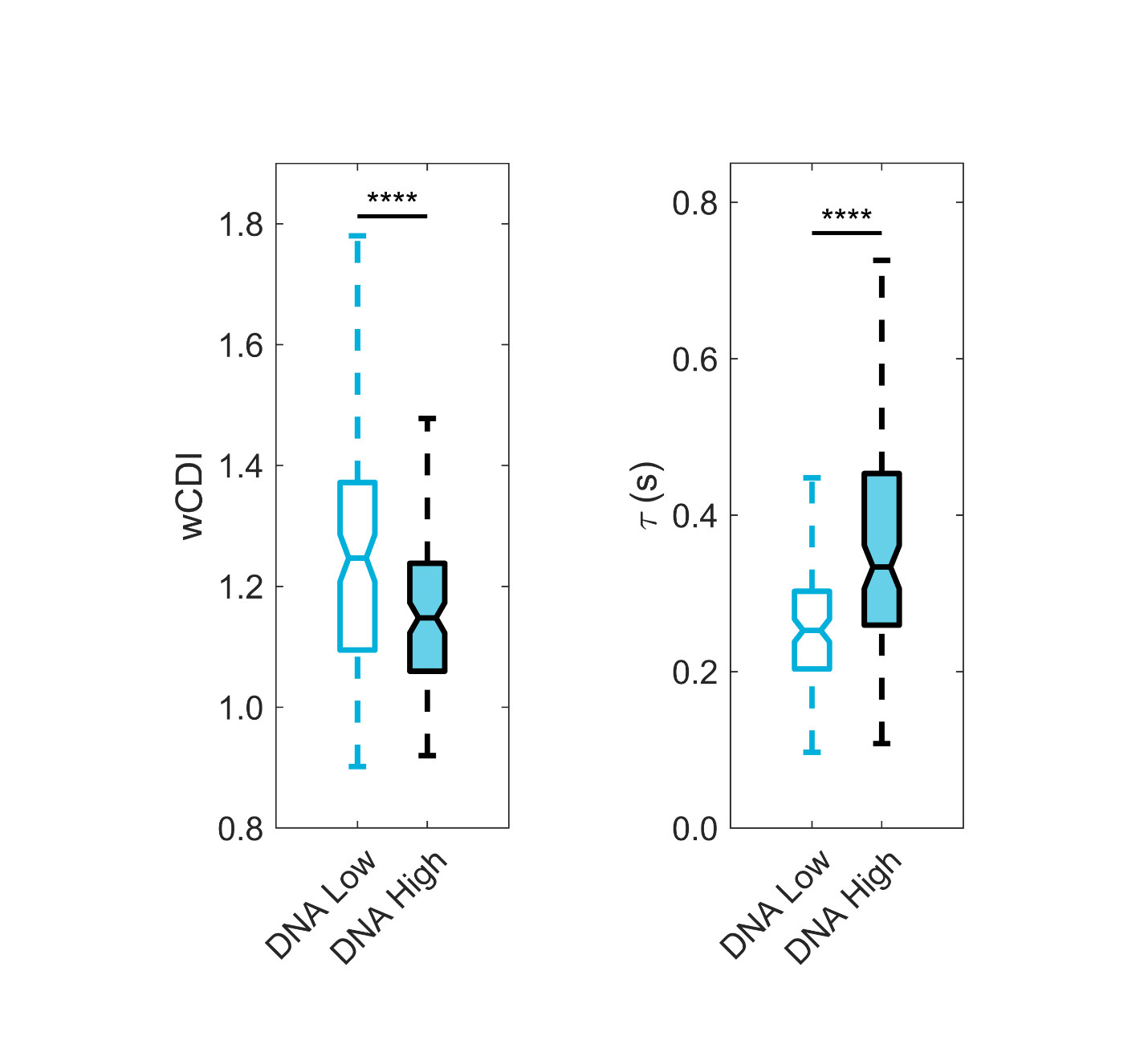

**Supplementary Figure S8:** Box plots of wCDI (left) and recovery times (right) for NB4 cells stained for DNA (Hoechst 33342) and sorted by DNA content. Notches represent 95% confidence intervals for the true median of each distribution. *n* $\geq$ 123 for each distribution across 3 different devices. *****p* < 0.0001, determined by two-sample Student’s *t*-tests.

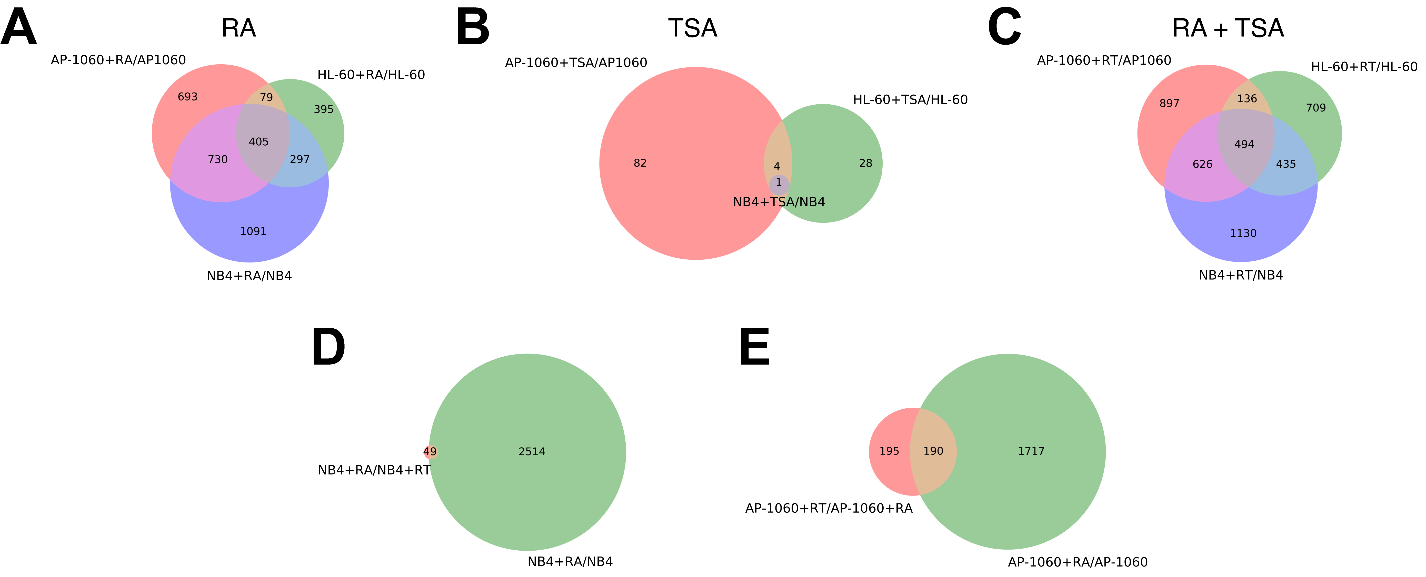
**Supplementary Figure S9:** Venn diagrams for differentially expressed genes identified via RNAseq. **A, B, C.** Untreated AP-1060, NB4, and HL-60 cells were compared to respective ATRA-treated samples (**A**), TSA-treated samples (**B**), and ATRA- and TSA-treated samples (**C**) per cell line. The sets of differentially expressed genes from each cell line-specific pairwise comparison were then compared. **D, E.** Comparisons to determine what effect TSA has on gene expression, in addition to or in interaction with ATRA. The set of differentially expressed genes for ATRA-treated vs. untreated samples was compared to that of ATRA- and TSA-treated vs. ATRA-treated samples for NB4 (**D**) and AP-1060 (**E**).

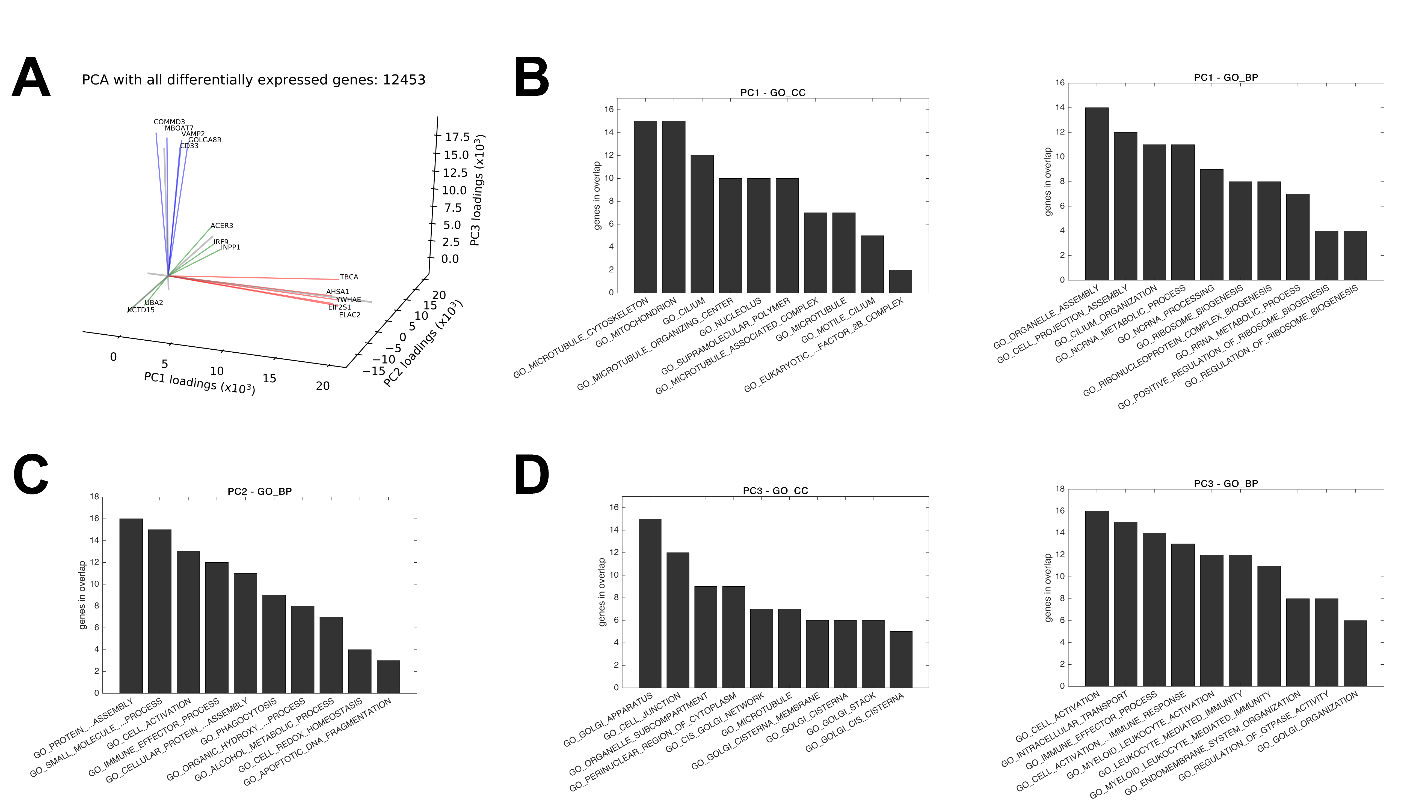
**Supplementary Figure S10:** Analysis of high-loading genes for 3-component principal component analysis for all 12,453 differentially expressed genes identified across all 66 pairwise comparisons between 12 triplicate samples. **A.** Biplot for top 5 genes of each principal component according to PC loadings. **B, C, D.** Gene set overlap between the top 100 highest-loading genes of PC1 (**B**), PC2 (**C**), and PC3 (**D**) and MSigDB C5 (gene ontology) gene sets for GO cellular component (GO_CC) and GO biological process (GO_BP). For PC2, only one GO cellular component gene set (GO_NUCLEAR_OUTER_MEMBRANE_ ENDOPLASMIC_RETICULUM_MEMBRANE_NETWORK, 13 overlapping genes) overlapped with input genes.

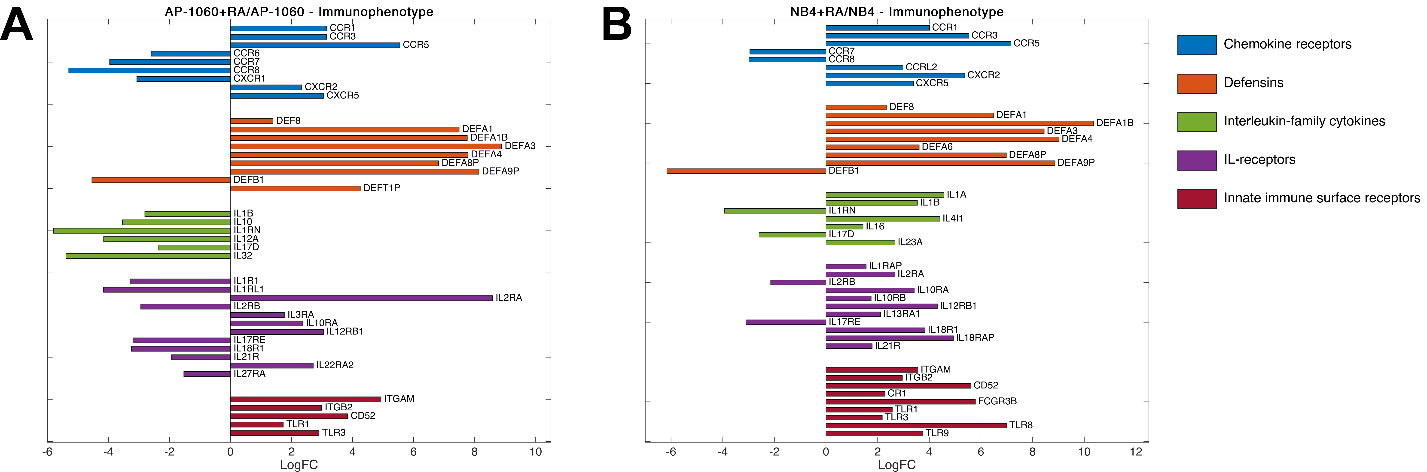
**Supplementary Figure S11: A, B.** Differentially expressed genes related to immune function and immunophenotyping identified in comparing ATRA-treated vs. untreated AP-1060 (**A**) and NB4 (**B**).

**References**

1.	DeBlois RW, Bean CP. Counting and sizing of submicron particles by the resistive pulse technique. Rev Sci Instrum. 1970;41(7):909–16.

2.	Saleh OA, Sohn LL. Quantitative sensing of nanoscale colloids using a microchip Coulter counter. Rev Sci Instrum. 2001;72(12):4449–51.
