## Supplementary figures and images for "Mechanical phenotyping of acute promyelocytic leukemia reveals unique biomechanical responses in retinoic acid-resistant populations"

### Supplementary Movie 1

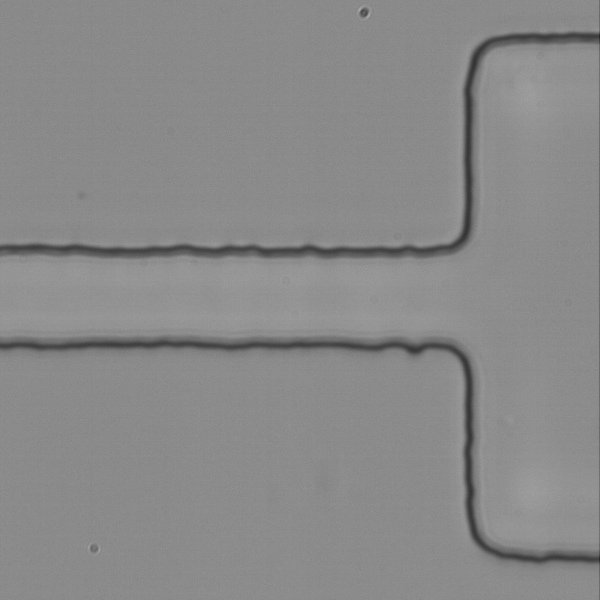

### Supplementary Movie 2

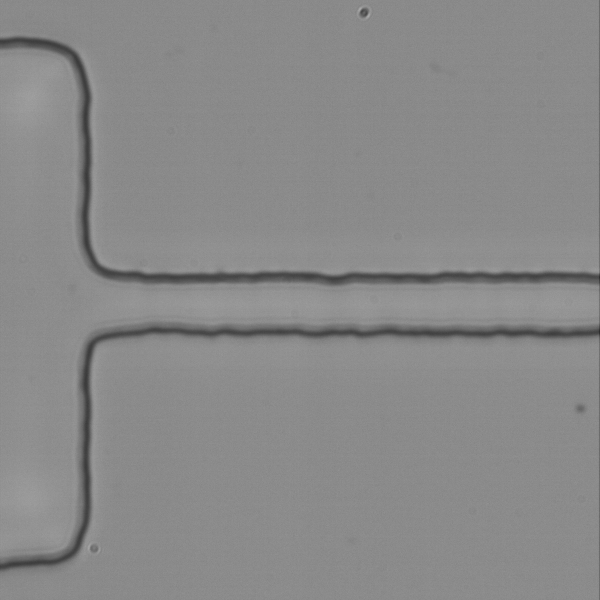

### Supplementary Movie 3

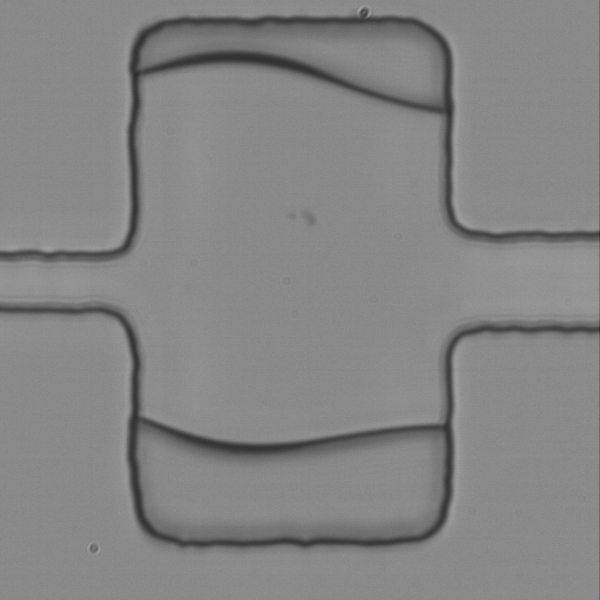
